# Dragonfly visual neurons selectively attend to features in naturalistic scenes

**DOI:** 10.1101/2020.09.14.297374

**Authors:** BJE Evans, JM Fabian, DC O’Carroll, SD Wiederman

**Affiliations:** The University of Adelaide; Lund University

## Abstract

Aerial predators, such as the dragonfly, determine the position and movement of their prey even when embedded in natural scenes. This task is likely supported by a group of optic lobe neurons with responses selective for moving targets of less than a few degrees. These Small Target Motion Detector (STMD) neurons are tuned to target velocity and show profound facilitation in responses to targets that move along continuous trajectories. When presented with a pair of targets, some STMDs competitively select one of the alternatives as if the other does not exist.

Here we describe intracellular responses of STMD neurons to the visual presentation of many potential alternatives within cluttered environments comprised of natural scenes. We vary both target contrast and the background scene, across a range of target and background velocities. We find that background motion affects STMD responses indirectly, via the competitive selection of background features. We find that robust target discrimination is limited to scenarios when the target velocity is matched to, or greater than, background velocity. Furthermore, STMD target discriminability is modified by background direction. Backgrounds that move in the neuron’s anti-preferred direction result in the least performance degradation.

**Significance Statement:** Biological brains solve the difficult problem of visually detecting and tracking moving features in cluttered environments. We investigated this neuronal processing by recording intracellularly from dragonfly visual neurons that encode the motion of small moving targets subtending less than a few degrees (e.g. prey and conspecifics). However, dragonflies live in a complex visual environment where background features may interfere with tracking by reducing target contrast or providing competitive cues. We find that selective attention towards features drives much of the neuronal response, with background clutter competing with target stimuli for selection. Moreover, the velocity of features is an important component in determining the winner in these competitive interactions.

## Introduction

An animal’s ability to detect prey, predators and mates within its environment can be essential to its survival. This task often involves the detection of small targets against highly cluttered backgrounds, where background features such as falling leaves, other animals, or even highly contrasting objects in the background scene may appear as potential targets. Numerous species have developed strategies for detecting small targets across a variety of sensory modalities, including auditory localization in bats (Arlettaz et al. 2001), the lateral line organ in squid (York et al. 2016) or visual cues as in insects (Nordstrom & O’Carroll 2006, Wiederman & O’Carroll 2011), archerfish (Schuster et al. 2006) and humans (Bravo & Farid 2001). Despite having relatively small brains and poor visual resolution (Horridge 1978, Land 1997), insects have solved the task of small target detection. This makes insects attractive models for investigating the neuronal computation underlying this complex behaviour.

Numerous strategies are employed by biological systems to improve target tracking in clutter. These include physical adaptations such as faster photoreceptors and more acute subregions of the eye (similar to the human fovea) thus improving spatial resolution (Hornstein et al. 2000, Burton & Laughlin 2003). Moreover, improved contrast sensitivity (Gonzalez-Bellido et al. 2011) can enable the detection of smaller and dimmer targets (Straw et al. 2006). Behaviourally, insects improve catch success by placing targets in such acute zones by directing their gaze (Olberg et al. 2007, Wardill et al. 2017). Dragonflies use predictive pursuit strategies (Mischiati, Lin et al., 2015) to intercept targets. While these adaptations improve target discriminability and pursuit success, how biological systems perform target-clutter discrimination is still poorly understood.

Neurons tuned to small moving targets are likely to underlie these behaviours and have been described from several flying species (Collett 1971; Collett & King 1975; O’Carroll 1993; Nordstrom & O’Carroll 2006; Geurten et al 2007; Keles, Frye 2017). In dragonflies, Small Target Motion Detector (STMD) neurons respond robustly to small targets (subtending less than 5 degrees) moving at any location within their receptive field (O’Carroll 1993). STMDs are sensitive to target contrast and are tuned to both the size and velocity of the target (O’Carroll & Wiederman, 2014).

CSTMD1 (Centrifugal Small Target Motion Detector 1), a well-studied STMD found in the dragonfly *Hemicordulia tau* (Geurten et al. 2007), has become an important model for investigating the neural mechanisms underlying target detection, even when embedded in natural images (Wiederman & O’Carroll 2011). CSTMD1 possesses both excitatory and inhibitory hemifields on opposite sides of the visual midline. CSTMD1 exhibits a predictive modulation of gain, which facilitates neuronal responses to targets moving on continuous trajectories in front of the prior path, whilst suppressing stimuli appearing at other locations (Wiederman, Fabian et al., 2017). Additionally, CSTMD1 exhibits selective attention, responding only to a single target, when presented with a pair of alternatives, completely ignoring the second target (Wiederman & O’Carroll, 2013, Lancer et al. 2019).

Past studies of STMD responses within visual clutter found that these neurons can respond robustly despite potential conflicting features of the background (Nordström et al., 2006, Wiederman & O’Carroll, 2011). However, the visual stimuli in these studies were limited to artificially generated backgrounds with low phase congruence (Nordström et al., 2006) or spatially constrained natural scenes that lacked relative motion cues (Wiederman & O’Carroll, 2011). While the ability of these neurons to respond to targets against the background even in the absence of relative motion is impressive, dragonflies operate in highly dynamic environments, where foreground (target) and background motion would vary dramatically, particularly during conspecific pursuit flights. How dragonfly STMD neurons respond to targets in natural scenes with varying degrees of relative motion remains unknown. Furthermore, the observation of selective attention in STMD neuronal responses raises the question of how such competitive processes interact with target-like background features embedded within the natural imagery, especially since prior studies have shown that relative motion is not a prerequisite for responses to such features (Nordström et al, 2006, Wiederman & O’Carroll, 2011).

Here we present STMD responses to moving natural images with superimposed, independently moving small targets to determine changes to target detection due to background clutter. These neurons exhibit reduced performance when targets are presented against visual clutter, particularly if the background speed is higher than that of the target. We show that this is not due to variable target contrast or inhibitory interactions from the background motion per se, but rather as the result of a competitive interplay between target and background features. Underlying these neuronal operations, we show the importance of precedence, with the first seen target more likely to be attended, even if it is inhibitory to the neuron. We also show the critical importance of velocity tuning, with CSTMD1 preferentially selecting faster background features that better match the velocity optimum over slower moving foreground target stimuli.

## Methods

### Electrophysiology

45 wild-caught, dragonflies (*Hemicordulia tau,* 39 male, 4 female*, Hemicordulia australiae,* 2 male) were immobilized with a 1:1 beeswax and rosin mixture and fixed to an articulated magnetic stand with the head tilted forward to access the posterior surface. A hole was cut above the brain to gain access to the lobula and lateral midbrain, but the preparation was otherwise left with the perineural sheath and overlying haemolymph sacs intact. We penetrated the sheath and recorded intracellularly using aluminosilicate micropipettes (OD=1.00, ID=0.58 mm), pulled on a Sutter Instruments P-97 puller and backfilled either with KCl (2M, electrode tip resistance typically 50-150 MΩ) or 4% Lucifer Yellow solution in 0.1M LiCl. Electrodes were placed in the medial portion of the lobula complex and stepped through the brain from posterior to anterior through the lobula complex, using a piezoelectric stepper (Marzhauser-Wetzlar PM-10). Intracellular responses were digitized at 5 kHz with a 16-bit A/D converter (National Instruments) for off-line analysis.

### Visual Stimuli

We presented stimuli on high definition LCD monitors (Asus ROG Swift PG279Q 165 Hz frame rate). The animal was placed 20 cm away and centred on the visual midline. Contrast stimuli were presented at screen centre (i.e. at the dragonfly’s visual midline) to minimize off-axis artefacts. The display projection was distorted using OpenGL to ensure each 1° onscreen was 1° from the animal’s perspective. The visual field was 104° (-52 to 52° azimuth) by 58.5° (21 to 80° elevation from equator).

For classification of neurons, we presented a sequence of stimuli: a gyrated, randomly generated texel pattern (1°), grey to black and grey to white full screen flicker (White 338 cd/m^2^, Black 0.5 cd/m^2^), moving edges (up, down, left and right, 25°/s), moving bars (2° width, up, down, left and right, 25°/s) and a square-wave grating pattern (0.025 cycles/°, 6.25Hz, up, down, left and right) and a small target (1.5°) moving at 80°/s. Neurons were identified based on their characteristic responses to these visual stimuli including their spike waveforms and unique receptive fields.

### Neuroanatomy

We identified a previously undescribed neuron, which we named ‘Binocular Small Target Motion Detector 2’ (BSTMD2) based on its binocular excitatory receptive fields and unique morphology revealed by intracellular labelling (Figure 1A, B, C). Following physiological characterisation, neurons were injected with Lucifer Yellow by passing 1nA negative current for 12 minutes. Brains were carefully dissected under phosphate buffered saline (PBS) and then fixed overnight in 4% paraformaldehyde in PBS at 4°C. To intensify the lucifer injection, brains were then rinsed (3x10 minutes) in PBS, before permeabilization in 80/20 DMSO/Methanol solution for 55 minutes and further rinsing (3x30 minutes) in PBS with 0.3% Triton X-100 (PBT). Brains were then preincubated in 5% normal goat serum in PBT for 3 hours at room temperature with gentle agitation followed by incubation in 1:50 dilution of biotinylated anti-lucifer yellow antibody (RRID: AB_2536191) in universal antibody dilution solution (Sigma Aldrich) for 3 days at 4°C with occasional gentle agitation. Brains were then rinsed (3x30 minutes) in PBT, followed by incubation with a 1:50 dilution of NeutraAvadin DyLight 633 for 3 days at 4°C. The samples were then rinsed in PBT, dehydrated through an ethanol series (70%, 90%, 100%, 100%), before clearing in methyl salicylate and mounting on a cavity slide in Permount for imaging. Following wholemount imaging of the neuron using mulitple Z series in a confocal microscope (Leica SP3, 10x Objective), image stacks were then stitched using the (Schindelin et al. 2012) and the neuron reconstructed in 3D using Neutube (Feng et al. 2015).

**Figure 1:**
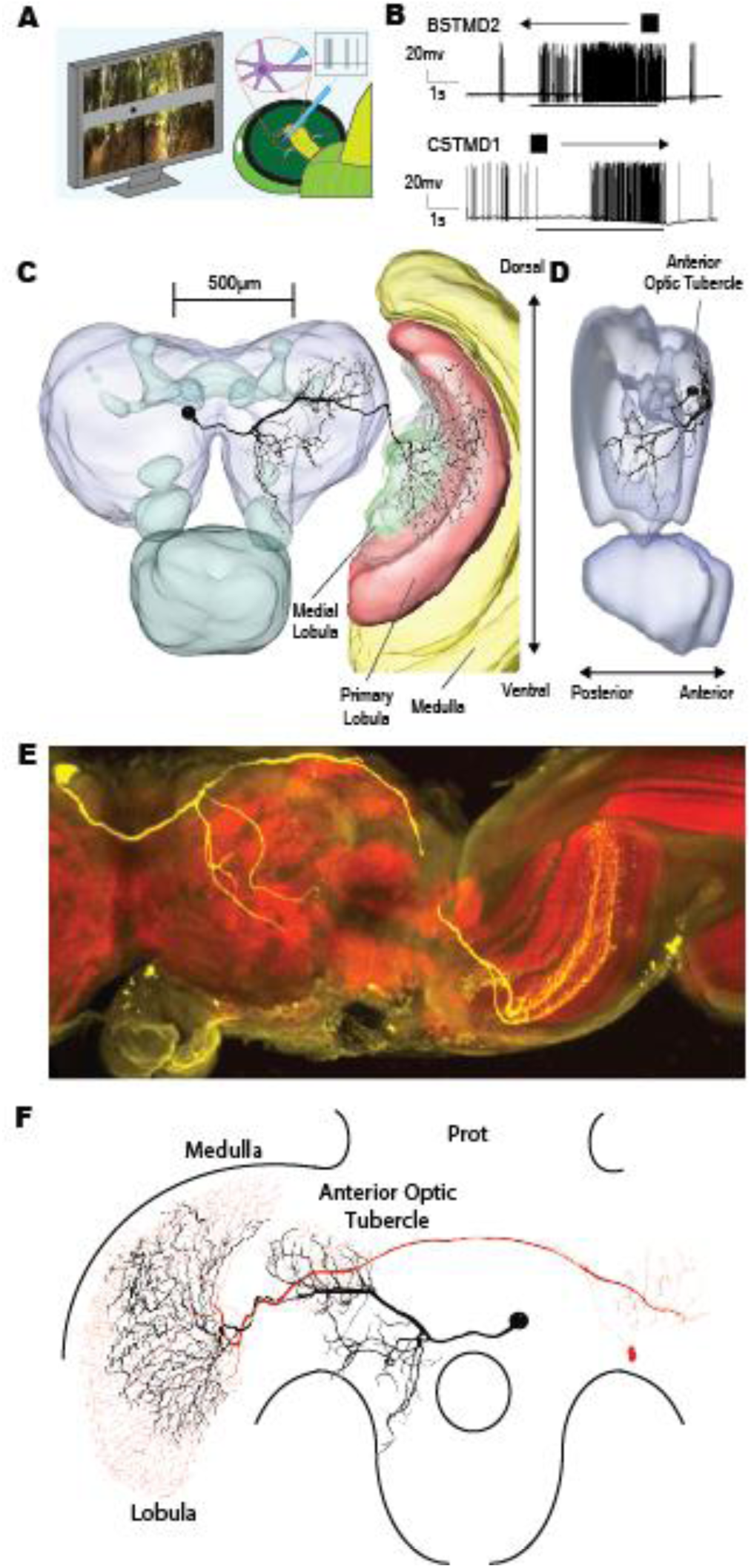
BSTMD 2 is a new large-field STMD responsive to leftward moving targets. Intracellular labelling of BSTMD2 reveals a centrifugal neuron arborizing widely in the medial and primary lobula. **A,** Illustration of intracellular recordings from the optic lobe whilst the dragonfly is presented with visual stimuli on an LCD display. **B,** Example raw traces of neuronal responses to a small square target drifting 80°/s through the receptive field of CSTMD1 (top) and BSTMD2 (bottom). Both neurons exhibit strong biphasic spiking responses (black bar indicates stimulus timing). **C, D,** BSTMD2 dye-fill superimposed over a Hemicordulia brain volume reconstruction. BSTMD2’s inputs arborize in the medial and primary lobula and has its cell body in the contralateral brain (compared to inputs). It also possesses a large arborisation in region corresponding to the Anterior Optic Tubercle (AOT). **E,** Microscope image of BSTMD2 dye-fill (yellow) and synapsin staining (red) of sectioned brain showing arborisation locations in lobula (see methods). **F,** CSTMD1 (red) and BSTMD2 (black) superimposed over simplified brain schematic. BSTMD2 AOT arborisations correspond with CSTMD1 arborisations.

In order to better visualise the lobula structures in which the neuron in Figure 1E arborized, we post-processed the brain in order to counterstain the synaptic neuropils using an anti-synapsin antibody (RRID:AB_528479). The coverslip was removed and the brain dissolved out of the Permount by immersion in xylene (3 hours at room temperature). Following rehydration through a descending ethanol series and resuspension in PBT, the brain was then prepared for vibratome sectioning, as described by (Heinze et al. 2012) by embedding in a gelatin-albumin mixture (4.8% gelatin and 12% ovalbumin in water), and allowed to set at RT before post-fixing overnight in 4% Paraformaldehyde at 4°C. 200µm horizontal sections were then cut on a Leica vibratome and collected in a 24-well plate before rinsing (6x20 minutes) in PBT. Sections were blocked using 5% normal goat serum in PBT, before incubation with anti-synapsin for 3 days at 4°C in the dark. Sections were then rinsed in PBT (6x20 minutes) before incubation with goat anti-mouse conjugated with CY3 at a dilution of 1:300, for 3 days at 4°C. The incubation solution also contained a 1:50 dilution of streptavidin conjugated CY5 in order to refresh fluorescence of the anti-lucifer antibody. Sections were then rinsed in PBT (6x20 minutes) before dehydration through an ethanol series (70%, 90%, 100%, 100%, each 20 minutes), before clearing in methyl salicylate and mounting with Permount. Images were then obtained using a Leica SP8 DLS confocal microscope with 20x oil immersion objective.

### Experimental Design and Statistical Analysis

We developed custom-written MATLAB scripts for spike counting and analysis. Curve fits used MATLAB’s in-built curve-fitting tools. All statistical tests were either paired t-tests or two-sided non-parametric tests (Kruskal Wallis test with Dunn’s multiple comparison correction). All p values are reported as raw numbers in text if significant differences exist (unmarked otherwise) or as < 0.0001 if sufficiently small. Box and whisker plots represent the 75^th^, 50^th^ and 25^th^ quartiles (lines) with raw data overlaid.

## Results

### Neuron Characteristics and Neuroanatomy

We first tested the ability of dragonfly visual neurons to discriminate targets in moving backgrounds by recording from STMD neurons with large receptive fields. Visual stimuli were presented on a display centred on the dragonfly’s midline (Figure 1A). We successfully obtained repeated intracellular recordings from two such neuron types, the well-characterised CSTMD1 (Geurten et al., 2007, Wiederman and O’Carroll 2013, Wiederman, Fabian et al., 2017) and a previously undescribed giant STMD neuron (Figure 1). Like the large-receptive field STMD neuron BSTMD1 previously described (Dunbier et al. 2012), this neuron gives excitatory responses to small moving targets in both visual hemispheres. We thus named this newly identified neuron ‘Binocular Small Target Motion Detector 2’ (BSTMD2). Individual examples of spiking activity in response to a moving target reveal robust responses in both CSTMD1 and BSTMD2 (Figure 1B).

Like CSTMD1, BSTMD2 is a centrifugal neuron, with two main optic lobe arborizations, one with bi-stratified dendrites in two adjacent layers in the outer part of the primary lobula, and the other a more compact arborization within a wedge-shaped deeper neuropil, the medial lobula (Figure 1E). The cell body is located on anterior brain surface, just contralateral to the esophageal foramen. The main axon is linked to the cell body and runs along the anterior surface of the lateral midbrain, where it connects with several regions of input or output (Figure 1C, D, E). One dense arborisation is found in an anterior structure which we putatively label as the anterior optic tubercle, at a similar location to the inputs of CSTMD1 (Figure 1F). Two second branches make sparse connections over a number of other more dorsal structures in the lateral midbrain.

### Direction Selectivity and receptive fields

BSTMD2 has a large binocular excitatory receptive field, with strong excitatory responses to small targets moving across either visual hemisphere. Unlike CSTMD1, BSTMD2 is also direction opponent exhibiting clear inhibition for rightward-moving targets (for the neuron recorded from the animal’s left optic lobe, Figure 2A, upper trace) and excitation for leftward-moving targets (Figure 2A, lower trace). Similar direction opponency has been previously observed in dragonfly STMDs (O’Carroll 1993) and more recently described in crab LCDC neurons, which exhibit some functional similarity to the dragonfly STMDs (Scarano et al. 2020). Like CSTMD1, BSTMD2 prefers small targets (Figure 2B, top) over elongated features (Figure 2B, bottom) but still shows modest responses.

**Figure 2:**
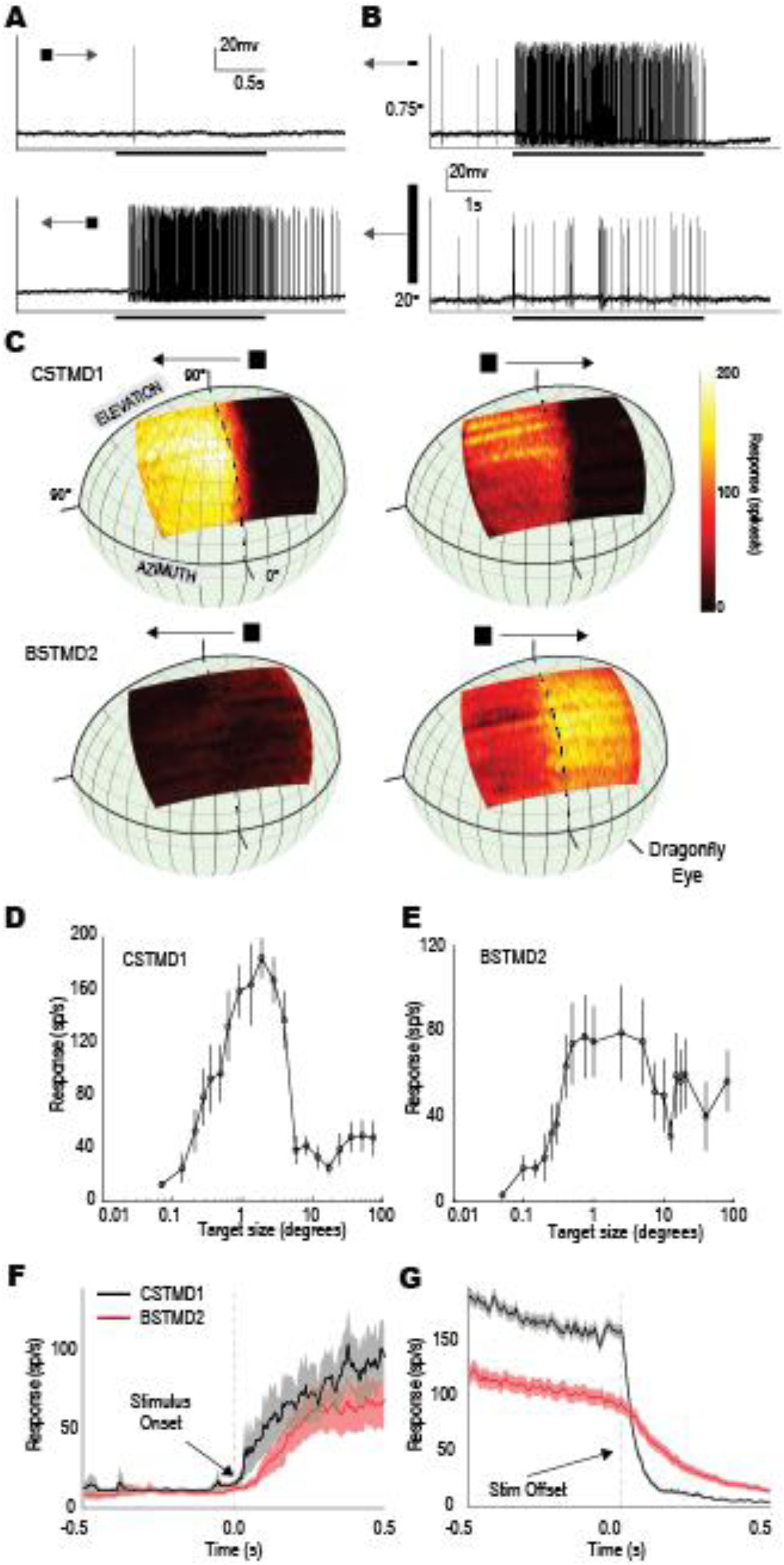
A, Raw spike traces showing BSTMD2s response to a moving target. BSTMD2 exhibits directional opponency, with inhibitory responses to rightward moving targets (top) and excitatory responses to leftward moving targets (bottom). **B,** Raw spike traces showing BSTMD2s preference to small targets (top) with reduced response to elongated bar features (bottom). **C,** Receptive fields of CSTMD1 and BSTMD2 as projected onto the dragonfly’s eye. Mean receptive field of CSTMD1 (n = 11) to a leftward (top left) and rightward (top right) moving small target and mean receptive field of BSTMD2 (n = 5) to a leftward (bottom left) and rightward (bottom right) small target **D,** Responses of CSTMD1 (n = 8) to bars of varying height (size orthogonal to stimulus motion). CSTMD1 exhibits height tuning optimal for targets of ∼1-3°. **E,** Target height tuning as in (D) for BSTMD2 (n = 6). BSTMD2 is more weakly size-tuned with the largest responses to targets measuring ∼0.5-4°. Unlike CSTMD1, BSTMD2 produces moderate responses more elongated features (bars) measuring >10° (target width of 1.5°). **F)** Plot showing the mean and standard error of stimulus onset (left, CSTMD1, n=8; BSTMD2, n=6) and stimulus offset (right, CSTMD1, n=21; BSTMD2, n=10). BSTMD2 exhibits slow onset and offset-time courses (100s of milliseconds to fully cease response to the target) compared to the faster kinetics of CSTMD1.

As previously described (Bolzon et al, 2009) CSTMD1 also responds to targets in both visual hemifields, but in this case with a clear distinction between excitatory and inhibitory responses at the central midline. Responses are excitatory in the right hemifield (contralateral to our recording site in the left lobula) or inhibitory in the left hemifield (Figure 1B, Figure 2C, top). This distinction holds in both hemifields for targets moving in either direction, although motion away from the midline (rightwards) elicits stronger overall responses in the right hemifield than motion in the opposite direction (Figure 2C, Right). As recently described, this weak direction selectivity results from differential facilitation of the response: CSTMD1 is initially weakly directional but exhibits increased direction selectivity as targets move along long trajectories, preferring targets moving upwards and away from the midline (Fabian et al. 2019). Similar facilitation may also explain the gradual build of more intense inhibition evident in receptive field maps for BSTMD2, as rightward moving targets commence motion from the left hand side of the display (Figure 2C lower left). The magnitude of excitation is also dependent on target location, with more robust activity to leftward motion as targets cross the midline into the left hemifield.

### Stimulus size selectivity

Both STMDs are length tuned (length here being defined orthogonal to the direction of motion) for moving targets presented with a constant width of 1.5°, parallel to motion. CSTMD1 prefers 1-3° (targets Figure 2D), while BSTMD2 is somewhat less selective, with a broader optimum for targets that are 1-4° (Figure 2E). CSTMD1 responses are weak for high contrast targets longer than 5°. In comparison, BSTMD2 does respond to these larger targets, although with weaker responses compared to those for smaller targets. In addition to their stimulus selectivity and receptive field shape, CSTMD1 and BSTMD2 have an interesting difference in their response kinetics: CSTMD1 shows less latency than BSTMD2, and also has a more sluggish offset in its response once targets disappear from the screen, compared to CSTMD1, outlasting it by several hundred milliseconds (Figure 2F,G).

### Selective Attention Across Hemifields in CSTMD1

To investigate the impact of background clutter on CSTMD1 requires analysis of how selective attention may operate across both hemifields. Given our analysis of the CSTMD1 and BSTMD2 receptive fields, these two neurons provide us with experimental access to two different target-detecting neurons, each with different excitatory and inhibitory drives: spatial location for CSTMD1 and opponent direction selectivity for BSTMD2. It has been previously shown that CSTMD1 exhibits ‘selective attention’, responding exclusively to one target when two alternatives are presented at the same time (Wiederman & O’Carroll 2013, Lancer et al. 2019). These experiments tested for selective attention to paired targets only within the excitatory hemifield (contralateral) of CSTMD1 (Figure 1C). Previous research, however, found that when two targets are presented simultaneously, one in each hemifield, average responses of CSTMD1 are strongly suppressed (Bolzon et al., 2009).

Is it possible that this inhibition is a result of selective attention in individual trials, leading to a suppressed average response when the target in the left (inhibitory) hemifield is attended? To test this, we reproduced a subset of the experiment conditions used in the earlier study (Bolzon et al., 2009) by drifting two alternative targets (1.5° x 1.5°) vertically through CSTMD1’s receptive field (Figure 3A), one in the inhibitory hemifield (T_1_) and one in the excitatory hemifield (T_2_). Each target was presented with a horizontal separation of 50°, well outside the small region of binocular overlap (∼10°) near the midline (Figure 3A). We also presented targets on both shorter (1s) or on longer (1.5 s) paths, either in the excitatory or inhibitory hemifield. Targets were either presented individually, or as a pair. In addition, we varied paired trials to test whether we could bias subsequent responses with a ‘primer’ stimulus located at one or other location. Hence either neither target was primed (i.e. T_1_ & T_2_ both had the shorter 1 s duration) or one of the targets was preceded by a 0.5 s primer at T_1_ or T_2_ respectively. Within the excitatory receptive field, such priming was recently shown to result in a bias of selection for the primed target (Lancer et al., 2019).

**Figure 3:**
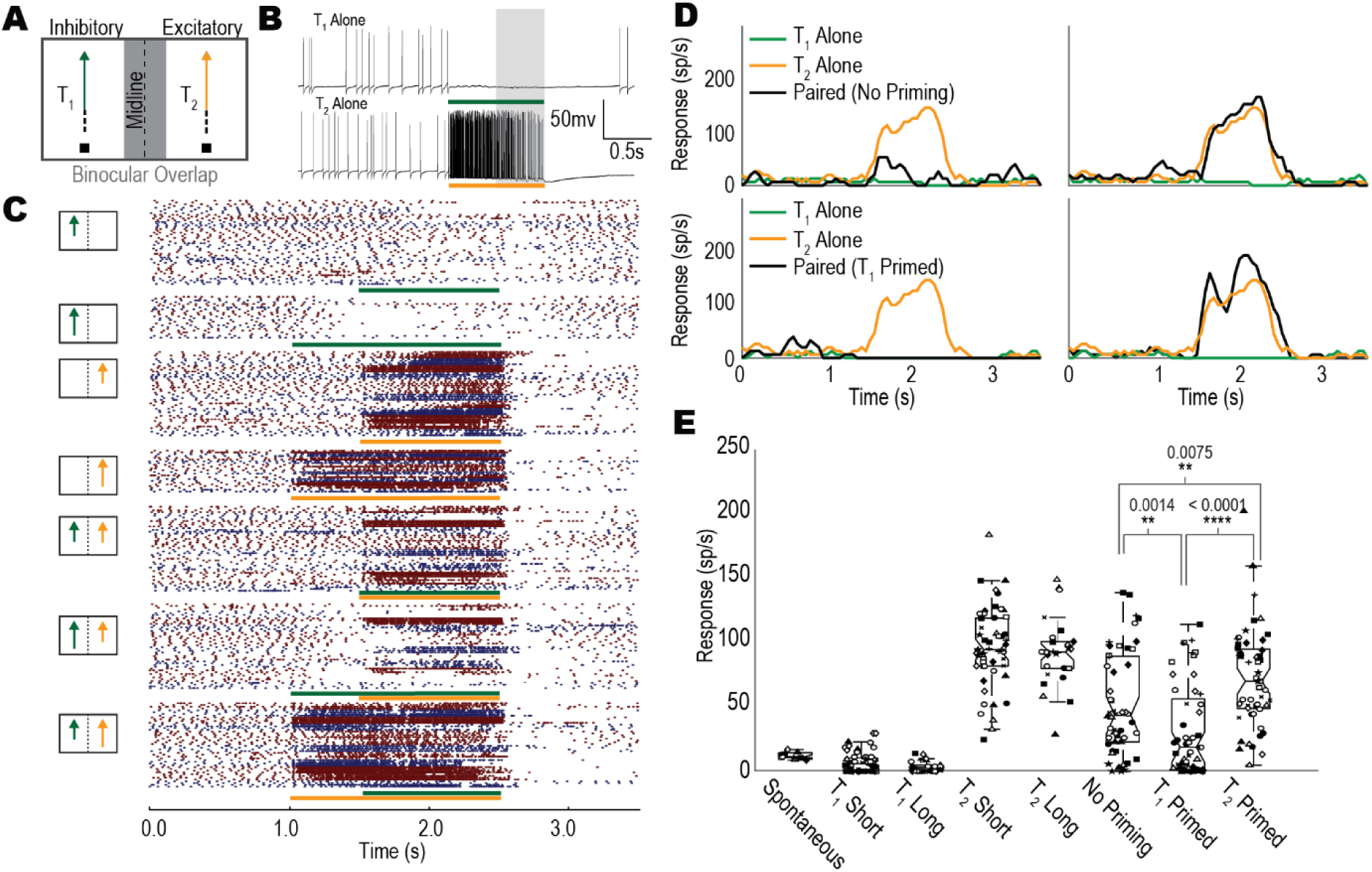
One target is selected when a pair of targets is presented on either side of the visual midline within CSTMD1’s receptive field. **A,** Targets are drifted vertically up the stimulus display, either individually or as a pair. T_1_ travels through the inhibitory region of CSTMD1’s receptive field. T_2_ moves through the excitatory receptive field. Targets are separated by 50° to avoid the 10° wide region of binocular overlap (grey region). **B,** Example neuronal traces of CSTMD1 responses to T_1_ (top) and T_2_ (bottom). T_1_ generates inhibition, T_2_ generates excitation. **C,** Raster plots over time (each point is a single spike) for single-target and paired target trials of CSTMD1. Green (T_1_) and orange (T_2_) bars indicate stimulus duration. Pictograms illustrate stimulus locations and length of the target trajectory (short, from 1.5 s to 2.5 s; long, from 1 s to 2.5 s). CSTMD1 trials from different dragonflies are separated by colour changes (n=8). T_1_-alone trials are inhibitory, reducing spike-rate to below spontaneous levels. T_2_-alone trials are excitatory, facilitating responses over longer trajectories. CSTMD1 responses to the paired target trials are either T_1_-like or T_2_-like, indicative of selective attention. This selection was biased by the presence of a preceding ‘primer’ target (inhibition when T_1_ primed, excitation when T_2_ primed). **D,** Depicts neuronal response (spikes/s) of different conditions taken from the same animal. In paired trials (no priming, green) we found cases where CSTMD1 responded similarly to the T_1_ alone case (top left) or the T_2_ alone (top right). Likewise, in T_1_ primed trials (light blue), we found responses which matched the T_1_ alone case (complete inhibition, bottom left) or the T_2_ alone case (excitation, bottom right). This emphasises the stochastic nature of the selection across hemispheres within a single animal. **E,** Each point represents the mean neuronal response over a 500 ms window (grey shaded region in B) prior to stimulus cessation (2-2.5s) for an individual trial (excluding spontaneous category). Different dragonflies (neurons) are marked with different symbols. Distributions of this data are presented in boxplots. In all paired-target trials there is significant variability with some trials exhibiting below-spontaneous activity (inhibition) and others strong activity (excitation). Priming T_1_ results in a shift of the mean response from the no-priming and T_2_-priming cases (p = 0.0086, p < 0.0001, n = 8, Kruskal Wallis multiple comparisons with Dunn post-hoc test).

Variability in responses between trials and dragonflies is evident from individual raster plots for all of these conditions from a number of recorded neurons (Figure 3C). Different dragonflies show different overall levels of spiking activity in the same identified neurons, likely due to small differences of stimulus location within the inhomogeneous receptive field. Each point in Figure 3C represents an individual spike, while the colour changes between red and blue indicate responses recorded from different neurons in sequential animals (n = 10). The stimulus conditions are shown in the pictograms to the left of each group of raster plots. The stimulus timings for each target are shown below the rasters (T_1_ green, T_2_ orange).

T_1_ presented alone generates strong inhibition, while T_2_ alone elicits strong excitation (Figure 3B), as would be expected from our receptive field analysis (Figure 2C). T_2_-only responses are stronger for the longer trajectory, consistent with the facilitatory effects observed in predictive gain modulation (Wiederman, Fabian et al. 2017). In T_1_-only trials presented in the inhibitory hemifield, inhibition is also increased for the longer trajectory stimuli. In paired target trials with no priming, we observed some trials in individual neurons that produced strong excitation while others produced strong inhibition. For the unprimed condition, individual dragonflies generally selected either T_1_ or T_2_, although we saw several examples of both excitatory and inhibitory responses to paired stimuli in a single animal, suggesting that these neurons can express competitive selection for targets across the two visual hemispheres (e.g. Figure 3D, top left and top right). For these un-primed, paired targets, the inhibitory target appeared to win more often, similar to the long-range inhibition previously observed in average responses (Bolzon et al., 2009).

In primed, pair trials, priming clearly influences which target is selected when subsequent stimuli are simultaneously presented in both hemifields. When the inhibitory hemifield was primed by a preceding target presented in only the inhibitory hemifield (T_1_ primed), responses during the paired stimuli were usually inhibited, (but not always, Figure 3D, bottom left, bottom right). Similarly, paired responses were more consistently excitatory when primed by stimuli confined to the excitatory hemifield (T_2_ primed).

To quantify these observations further, we counted spikes in a ‘late’ response window, 0.5 s in duration but commencing 0.5 s after the appearance of the second target (green shaded region in Figure 3B). This late window was selected so that even the shorter path (unprimed) conditions still provide time for the response to facilitate (Nordström et al., 2011). We aggregated this late response activity across trials and neurons for each of the conditions (Figure 3E). We found significant differences between all three two-target conditions, with T_2_ (excitatory) priming eliciting stronger subsequent responses (p = 0.0075, Kruskal-Wallis with multiple comparisons) than the unprimed case, while T_1_ priming leads to weaker responses (p=0.0014, Kruskal-Wallis with multiple comparisons).

Our aggregated plots (Figure 3E) also show the late window response from individual trials using varying symbols for different individual dragonflies. This confirms that the three primed paired conditions show trial-to-trial responses generally equivalent in magnitude to either a single excitatory or inhibitory target, thus tending to cluster bimodally between individual neurons or within (e.g. neurons ▴, □) In many trials, the T_1_ priming established strong inhibition, however this biasing was not always effective. In some trials, following strong inhibition by the primer target, responses in the late window then switched to the excitatory target, in either all trials (e.g. neuron △) or in a subset of trials (e.g. neurons □, ○).

Hence, while our results generally agree with the earlier work (Bolzon et al. 2009) in that we observed consistent inhibition when paired trials are averaged across many trials, we do also reveal trial-by-trial variability consistent with a competitive selective attention mechanism (Wiederman & O’Carroll 2013) whereby either visual hemisphere may sometimes be the ‘winner’ for attention in paired trials.

### Selective Attention across directions in BSTMD2

We also tested for selective attention in BSTMD2 using an equivalent set of conditions as used in CSTMD1. In this case we exploit the opponent direction selectivity of BSTMD2, rather than the spatial location of the stimulus as the competitive inhibitory or excitatory drive (Figure 4A). We drifted one target in the inhibitory direction, left to right, (T_1_) and the second target in the excitatory direction, right to left (T_2_). Figure 4B shows individual examples of BSTMD2 responses to targets drifted in either direction. As before, each target was presented for either 1 s (unprimed) or 1.5 s (primed). The targets were separated vertically by 10° and the positions pseudo-randomly swapped (i.e. either T_1_ or T_2_ at the top) to avoid bias from any receptive field inhomogeneities. In longer recordings, the location of the experiment within the receptive field was changed to avoid the effects of habituation.

**Figure 4:**
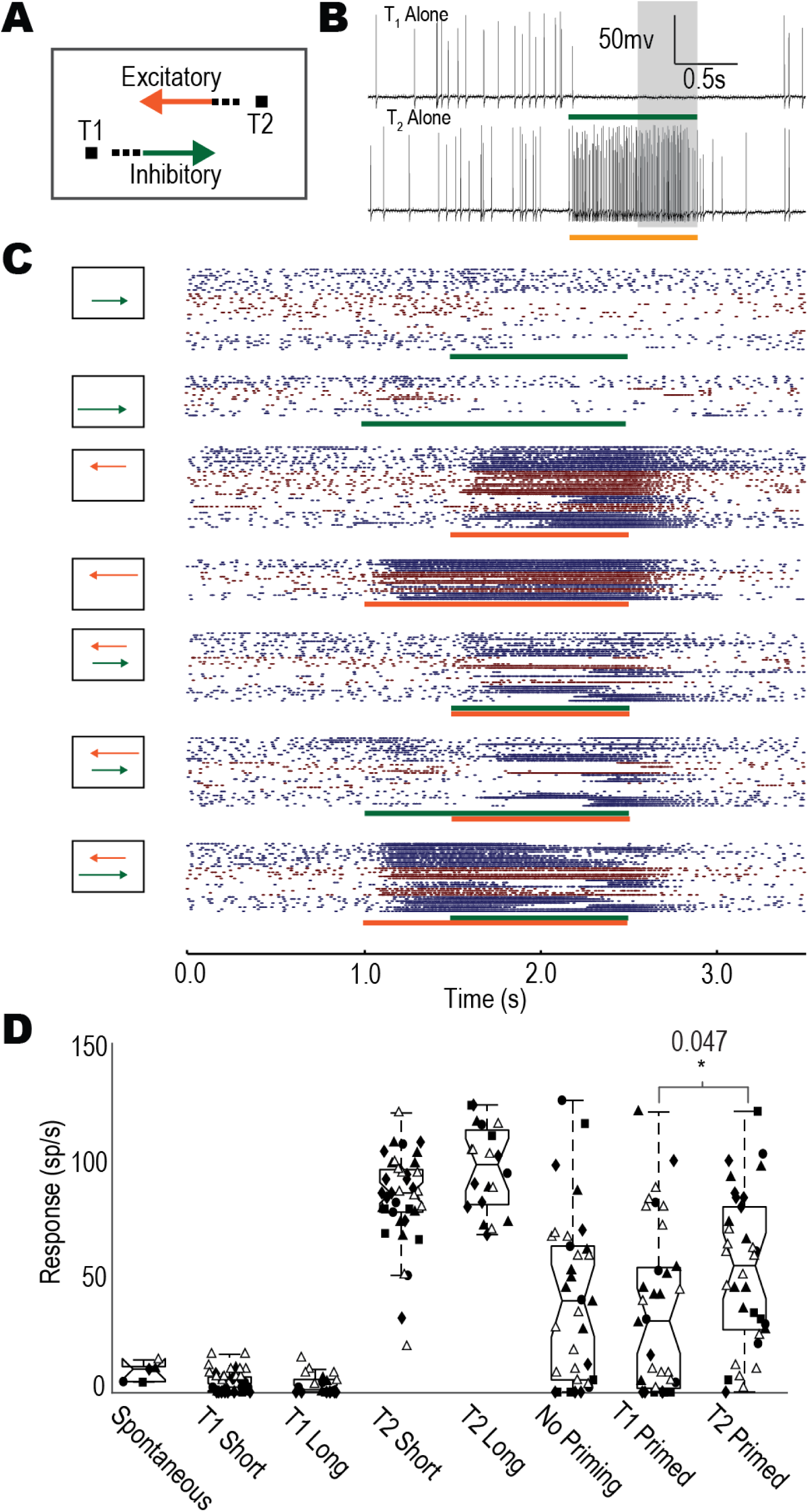
One target is selected when a pair of targets is presented moving in opposing directions within BSTMD2’s receptive field **A**, Targets are drifted horizontally across the stimulus display, either individually or as a pair. T_1_ travels in the inhibitory direction of BSTMD2’s receptive field. T_2_ moves in the excitatory direction. Targets are separated by 10° to limit surround antagonistic interactions (underlying size tuning). **B**, Example neuronal traces of BSTMD2 to T_1_ (top) and T_2_ (bottom). T_1_ generates inhibition, T_2_ generates excitation. **C**, Raster plots (each point is a single spike) for single-target and paired target trials of BSTMD2 (blue and red bars indicated stimulus time, T_1_ & T_2_ respectively, pictograms indicate stimulus locations, different animals are separated by colour, n=6). Paired target trials exhibit both T_1_-like and T_2_-like responses indicative of a selective attention mechanism. Primed trials shift these responses to the primed target (inhibition T_1_ primed, excitation T_2_ primed). **D**, Boxplots showing different conditions shown in (**C**). Each point represents the mean response over a window of 500ms (grey shaded region in **B**) just prior to stimulus cessation (i.e. 2.0-2.5s) for an individual trial (excluding spontaneous estimates which are averaged across neurons). Different animals are marked with different symbols. No statistical difference seen between primed and unprimed trials. T_1_ primed and T_2_ primed trials are significantly different (p = 0.034, Kruskal-Wallis with multiple comparisons test) indicating priming is effective in BSTMD2.

Figure 4C shows raster plots for individual trials in each of the seven conditions, equivalent to Figure 3. Single target trials in BSTMD2 elicited either strong inhibition or excitation, but in BSTMD1, this now depended on the attended target’s direction rather than its location in the visual world as we saw in CSTMD1. In the unprimed paired-target trials we observed a combination of both strongly excited and strongly inhibited responses. Interestingly, apparent switching from attending to one target or the other was far more prevalent in BSTMD2, with many examples of the responses switching from strong excitation to strong inhibition during a trial (and vice versa). The amount of excitation or inhibition at any instant broadly matched the response of the equivalent single-target trials (either T_1_ or T_2_ alone, Figure 4D). Primed target trials, where the primer was inhibitory (T_1_) resulted in more overall inhibition during the paired presentation. Similarly, an excitatory primer (T_2_) typically resulted in more overall excitation (Figure 4C).

Figure 4D shows the responses from the late window as described earlier for Figure 3E. We again found a significant difference in overall response between the T_1_-primed and T_2_-primed case (p = 0.034, n = 6, Kruskal-Wallis with multiple comparisons test). In unprimed paired-target trials there were numerous examples of individual cells (⧫, ▴, ▪, ●, △) that showed both excitatory and inhibitory responses. There were also many examples where the response was intermediate, though since we are here averaging the response over 0.5s, these can be attributed to mid-trial attention switches, which were common (see raster plots, Figure 4C). Priming appeared less effective than in CSTMD1. In the T_1_-primed trials four neurons (⧫, ▴, ●, △) exhibited significant response variability (both excitatory and inhibitory responses). Likewise, four T_2_-primed neurons (⧫, ▴, ▪, ●, △) were highly variable.

Our data shows that selective attention is not unique to CSTMD1 (and the BSTMD1 neuron from previous studies). Moreover, the winner of a winner-takes-all process can be expressed by stimuli that are inhibitory (via directional opponency) in the recorded neuron. Combined with the results from CSTMD1 (Figure 3), these results are indicative of a global selection system operating across hemifields and expressed in numerous neurons simultaneously. It is apparent that this competition can occur across hemifields, across directions and include inhibitory stimuli. These findings are vital to understanding how STMDs might react to complex cluttered scenes (presented below) where target-like features of the background may compete for selection.

### Minimizing Contrast Variation in Natural Images

Dragonfly STMDs are primarily responsive to dark targets moving against bright backgrounds (Wiederman et al. 2013). However, even a dark target varies in local contrast as it crosses light and dark regions in the background of natural scenes. This variation will modulate STMD responses over time due to the neuron’s strong sensitivity to contrast (O’Carroll & Wiederman 2014). Such changes will not only be due to ‘local’ intensity variation in both space and time (i.e. contrast), but also from longer-term adaptive (Wiederman et al., 2008) and self-facilitatory effects from prior history (Wiederman, Fabian et al., 2017). In addition, STMDs are influenced by spatial inhibitory interactions that underlie their size selectivity.

To investigate the effect of background clutter on STMD responses we developed a stimulus (similar to Nicholas et al., 2018) that limited these local interaction effects between target and background (Figure 5). Our stimulus consisted of one of three natural images (Figure 5A) with a 1.5x1.5° target moving at 25°/s horizontally across the field of view (CSTMD1 responds robustly to rightwards moving targets in its excitatory hemifield). The natural image was presented 1 s prior to the appearance of the target to avoid onset transients created by the appearance of the background. To avoid the effects of local contrast variation, we superimposed a grey strip over the top of the natural image along the intended trajectory of the target. The grey strip was 11° in height with a further 2° on either side linearly ‘fading’ to invisible via modulation of the alpha channel. The grey strip ensured that 1) the local target contrast was constant, 2) there was no texture varying the degree of local temporal adaptation and 3) short-range spatial inhibitory interactions underlying size selectivity (Bolzon et al. 2009) were minimized.

**Figure 5:**
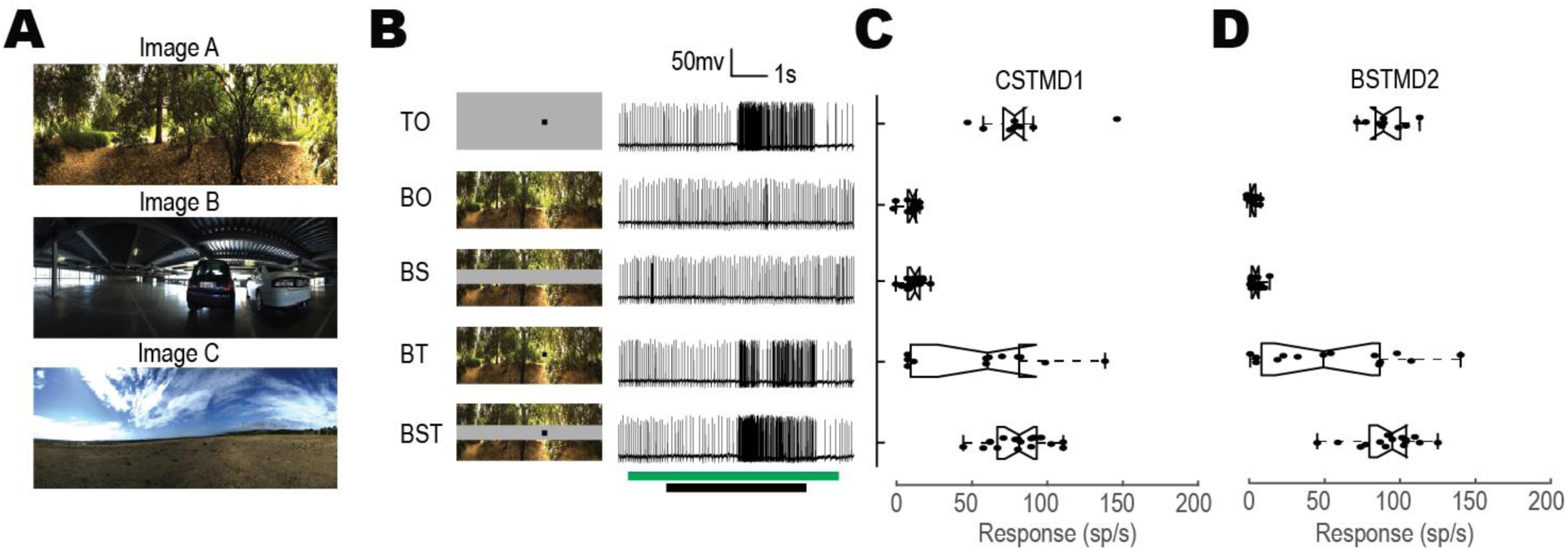
Background effects of contrast variation and size selectivity on CSTMD1 and BSTMD2 responses are limited by the inclusion of a grey strip. **A,** Three stationary natural images were presented to the dragonfly, chosen for their differing degrees of clutter. **B**, Each stimulus set was composed of variants, target-alone (TO), background-alone (BO), background with strip (BS), background with target (BT) and background with target and strip (BST). Example CSTMD1 responses are shown to each variant of one image in the set. Moving targets (during black stimulus bar) elicit robust responses compared to the presence of the static background (green stimulus bar). Variation induced by a target moving across the background (BT) is decreased by the presence of the grey strip (BST). **C,** Comparison of the five conditions in CSTMD1 (n=2). Each point represents the mean spike-rate in a 500 ms analysis window (corresponding to the strongest part of the receptive field), with aggregate data shown in boxplots. Variation on the target response induced by the variable background contrast (BT) are reduced by the addition of the grey strip (BST) with responses shifted towards those observed with the target alone (TO) condition. **D,** This method of deconfounding local contrast and surround antagonism is also effective in BSTMD2 (n=3).

We tested the efficacy of the grey strip in a small sample of neurons for one of five conditions (Figure 5B): moving target-only (TO), stationary background-only (BO), stationary background with grey strip (BS), stationary background with moving target (BT) and stationary background with grey strip and moving target (BST). As expected, both CSMTD1 (Figure 5C) and BSTMD2 (Figure 5D) show more varied responses during BT trials, as target contrast varies as it moves across different background features. However, the inclusion of the grey strip (BST) nullified this variation, returning the responses to a range equivalent to the TO trials.

To quantify this, we first calculated the TO mean response across all trials. Then for each image separately, we segmented the mean TO, individual BT and individual BST trials into 5 ms bins. We determined the difference for each bin between mean TO and BT trials, as well as mean TO and BST trials. We performed a one sample t-test to determine whether these differences were non-zero. We found that the error between TO and BT were non-zero for both CSTMD1 (p = 0.011) and BSTMD2 (p = 0.0054). However, there was no difference between the TO and BST for either CSTMD1 or BSTMD2 (p > 0.4). Thus, the grey strip paradigm was sufficient to remove contrast variations caused by changes in the background luminance.

### Target Tracking with Background Motion

We previously showed that CSTMD1 can respond robustly to small targets when they move at the same velocity as the background (Wiederman and O’Carroll, 2011). What if the target moves independently of the background, inducing relative motion cues?

We tested this using the same three panoramic images (Figure 6A), with each moving either leftwards or rightwards at 15°/s. The starting position (phase) of the backgrounds was also varied (0° and 180°) to reduce phase-dependent effects. The target (1.5x1.5°) moved at 25°/s, always in the preferred direction (rightwards for CSTMD1 or leftwards for BSTMD2, Figure 6A). The target speed was chosen to elicit an approximately half-maximal response compared to an optimally chosen velocity (Dunbier et al., 2012). The background speed was set to be lower than that of the target to ensure relative motion. In all cases, the background was presented 1 s prior to the start of the target which then moved across the entire stimulus monitor. We used five different stimulus conditions (Figure 6B): moving target alone (TO), leftward moving background with strip (BSL), rightward moving background with strip (BSR), moving target with leftward moving background and strip (BSLT) and moving target with rightward moving background and strip (BSRT). The background direction and timing are indicated by green arrows and green stimulus bars. The target direction and timing are indicated by black arrows and black stimulus bars (Figure 6B).

**Figure 6:**
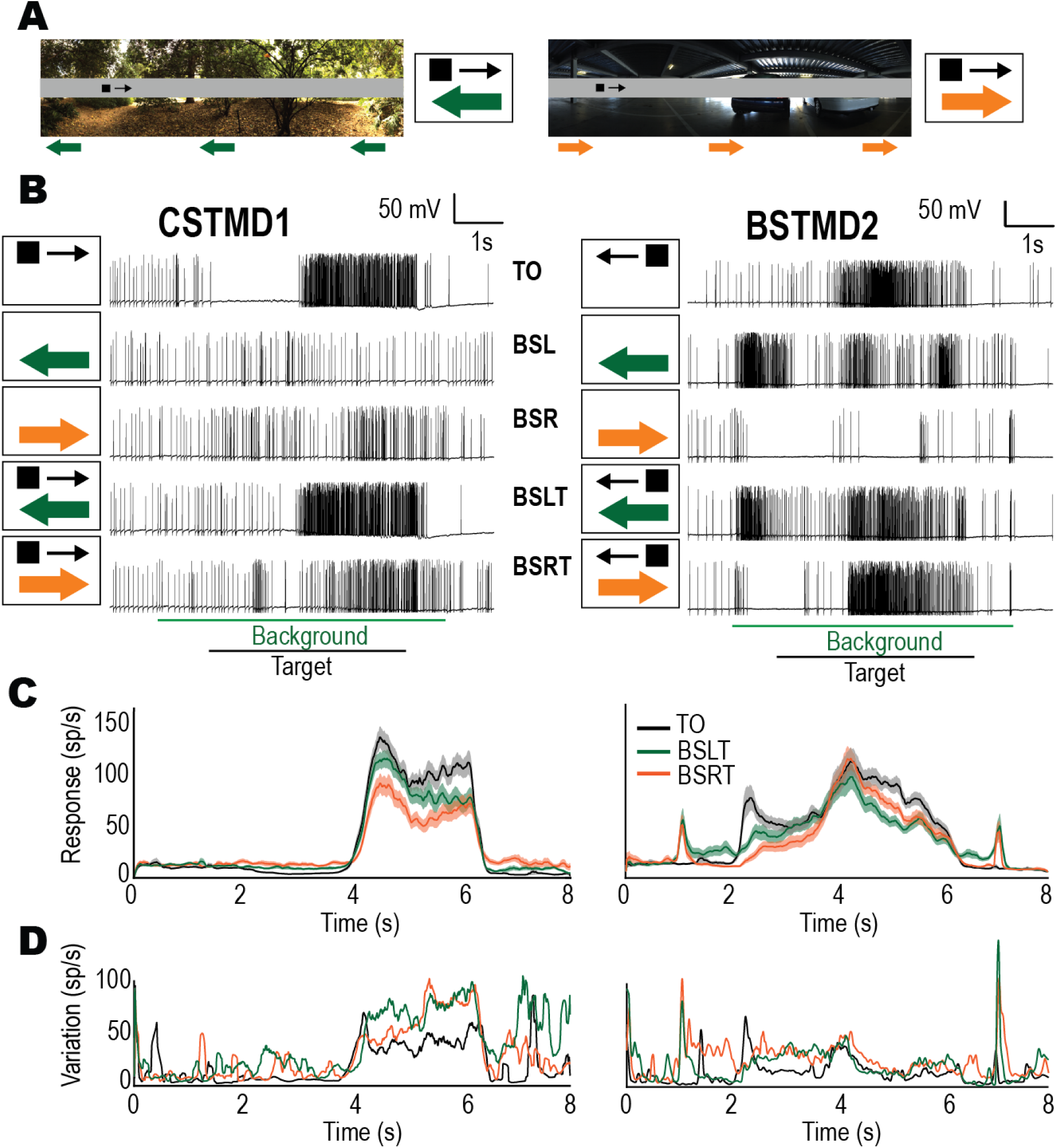
Responses of both CSTMD1 (n=12) and BSTMD2 (n=6) are reduced by backgrounds moving in either direction. A, Example of stimuli with either a leftwards or rightwards moving background (green arrows) and rightwards moving target (black arrow). Corresponding pictograms are illustrated. B, Example raw traces from CSTMD1 and BSTMD2 when presented with: target alone (TO), leftward moving background alone (BSL), rightward moving background alone (BSR), target with leftward background (BSLT) and target with rightward background (BSRT). Background features can both elicit activity as well as decrease responses. C, Mean spike-rate and standard error (shaded regions) of neurons (CSTMD1 left, BSTMD2 right) to TO, BSLT and BSRT. Targets with leftward or rightward backgrounds result in robust, but reduced spike-rates. D, Metric of variation (variance/mean) calculated from plots in (C). In CSTMD1 and BSTMD2 the variation increases for the two moving backgrounds, revealing that proportionally, variance increases with respect to the mean spike activity, a result incongruent with a constant inhibitory drive from a wide field motion pathway.

Individual examples of CSTMD1 responses to these five stimulus conditions (Figure 6B, left) show that while the background-only trials (BSL, BSR) generate only weak responses (either excitation or inhibition), the introduction of clutter in the background-with-target trials (BSLT, BSRT) significantly changed the responses from the equivalent TO trial (note the lack of inhibition in the BSLT trial and the weaker excitation in the BSRT trial, Figure 6B, left).

Additionally, when background and target were presented together, the overall average response to the target across trials was reduced. This reduction was larger when the background moved in a rightward direction (BSRT) compared to the leftward direction (BSLT).

Individual examples from BSTMD2 neurons (Figure 6B, right) show the profound effect that the addition of the background has on responses (BSLT, BSRT) compared with the target only (TO) equivalent. This is in part due to the increased sensitivity of BSTMD2 to rotated panoramas, which we attribute to its broader size-selectivity (preferring small targets but still giving intermediate responses to larger features, see Figure 2E). The effect of background motion in the background-only trials (BSL, BSR) in BSTMD2 are far more pronounced, with the leftward moving background eliciting more robust responses than its CSTMD1 counterpart, while the rightward moving background induced significant, sustained inhibition. This direction opponency yields a particularly interesting response for the BSRT trial shown in Figure 6B (right), dominated initially by the onset of inhibition due to the background, which persists for more than a second after the target appears before apparently switching attention to the target after which the response ‘breaks through’ to subsequent strong excitation.

We aggregated these responses across cells, by separating each trial into 5 ms bins and calculating the mean and standard error across all trials (including images and images phase) for each of the five conditions. Three of these conditions (TO, BSLT, BSRT) are shown in Figure 6C (CSTMD1 left, BSTMD2 right). In both background-with-target conditions (BSLT green line, BSRT orange line) responses were reduced compared to the target-only (TO) condition. In CSTMD1, the BSRT condition exhibited the stronger reduction in response. This is surprising given that leftward moving features more strongly stimulate the inhibitory hemisphere of CSTMD1, whereas rightward moving features more strongly excite the excitatory hemisphere of CSTMD1. In the first half of the target’s trajectory (i.e. whilst the target is within the inhibitory hemifield of CSTMD1, 2-4 s), the order of response strength was reversed with TO showing the most inhibition, followed by BSLT and finally BSRT.

Similar reductions in response were observed in BSTMD2. Prior to the target’s appearance and after its disappearance in BSLT trials, (at around 1-2 s and 6-7 s), responses were higher than the TO cases due to activity induced by background motion. The response reductions seen in BSTMD2 were not as consistent as in CSTMD1, although they showed similarly pronounced differences between hemifields. In the first half of the target trajectory (∼2-4 s), BSRT trials showed the weakest responses (likely due to the inhibitory rightwards moving background), while in the second half of the target trajectory (4-6 s), BSLT showed weaker responses.

Are the reductions in mean response to the target when seen within a background (BSLT, BSRT) due to a consistent inhibitory effect of the background pattern? In dragonflies, wide-field motion sensitive Lobula Tangential Cells (LTCs) give sustained velocity dependent responses to translated natural images (Evans et al. 2019). Could inhibitory input from this widefield, optic-flow system be responsible for the response changes we see in the background with target (B+T) cases (BSLT, BSRT)? To test this, we calculated the Fano Factor (Figure 6D), a metric of variation defined as the variance divided by the mean response. In both CSTMD1 (Figure 6D left) and BSTMD2 (Figure 6D right) the total amount of variation is higher in the background-with-target (BSLT, BSRT) trials. In CSTMD1 this increase in variation occurs in both the inhibitory (2-4 s) and excitatory (4-6 s) hemispheres. In BSTMD2 this additional variation occurs over the majority of the time the target is present except when the target is in the centre of the receptive field.

The increased variation is more indicative of a selection-like process than consistent inhibition. Were the response reductions observed caused by a wide-field motion-sensitive inhibitory feedback, we would expect the variation in response to reduce proportionally with the reduction in mean response, resulting in a relatively constant variation metric. If instead, the target system is selecting from either background features (weakly excitatory or inhibitory) or the target, we would expect a larger variation overall, even with a modest reduction in the mean response. To further illustrate these inter-trial variations, we generated raster plots for each of the individual trials separated by condition (Figure 7). For CSTMD1, we observed robust responses when the target was presented alone (Figure 7A). A target generates inhibition in the left hemifield (2-4 s,) and excitation in the right hemifield (4-6 s). When the background was presented alone (BSL, BSR), the responses were weak and intermittent (Figure 7B). In the B+T trials (BSLT, BSRT), the responses to the target is still discernible in most trials but is often weaker or less consistent (Figure 7C).

**Figure 7:**
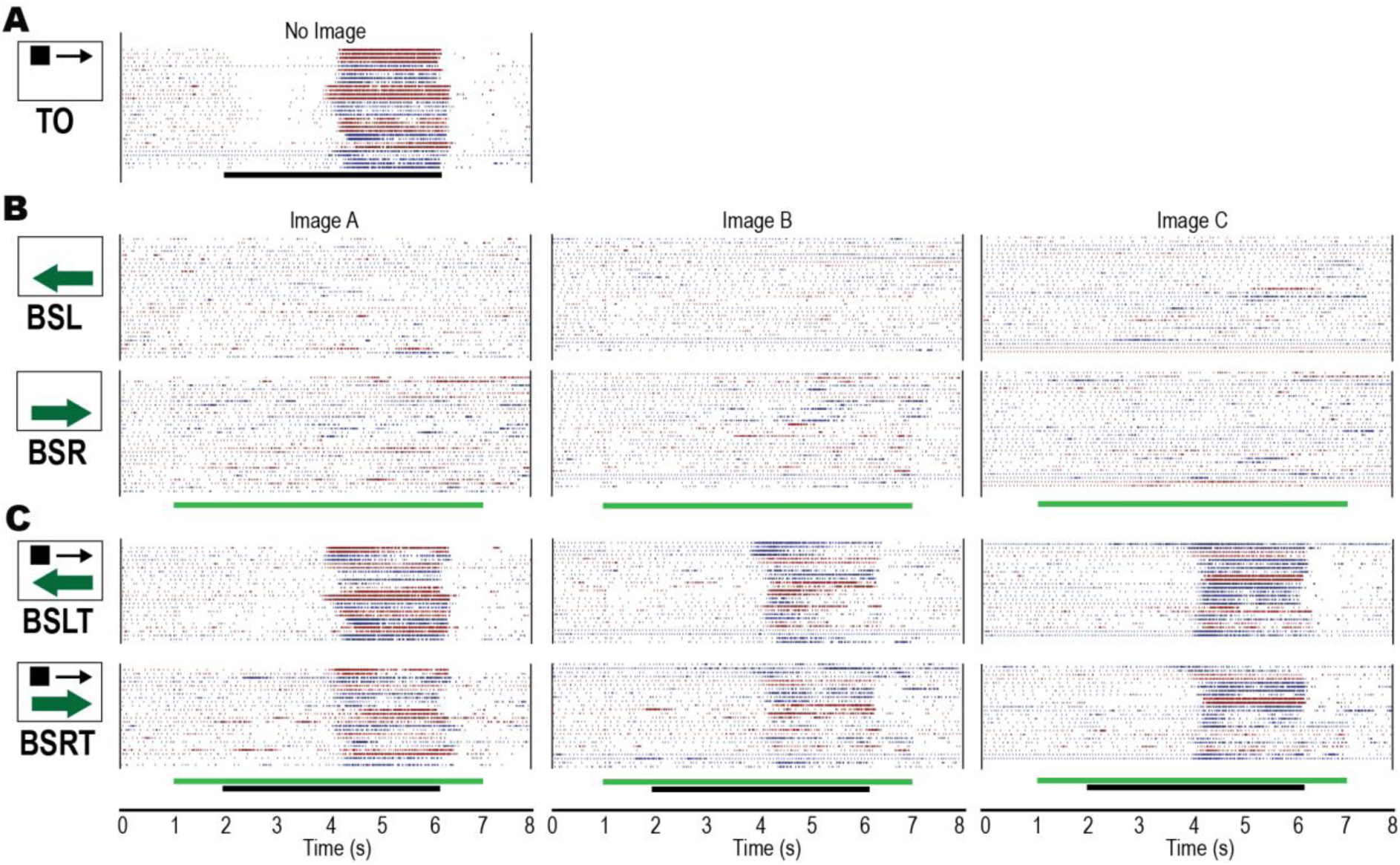
CSTMD1 raster plots reveal that moving backgrounds (15°/s) cause target responses to become less sustained and consistent. **A**, Target only responses (15°/s) (TO). Black stimulus bar indicates time target is moved on the display **B**, Responses to leftward moving backgrounds (BSL) and rightward moving backgrounds (BSR). Green stimulus bar indicates time background is moved on the display. The moving backgrounds result in intermittent, weak responses. **C**, Backgrounds moving in either direction, with a target moving in the preferred (BSLT, BSRT), cause a reduction in target response compared to target alone responses.

From the raster plots, it is also possible to identify where this response reduction occurs. Most individual responses retained their most robust response to the target as it approached the midline of the animal (just after 4 s) with much of the reduced response occurring when the target was located at a more peripheral location (5-6 s). It is also clear from the raster plots that BSRT trials have a greater target response reduction than those of BSLT. If this reduction is caused via a selection process, this indicates that these response reductions are most likely caused by low-salience distractors within the excitatory receptive field generating weakly excitatory responses. This matches previous findings where low contrast targets which appeared prior to a high contrast distractor often won the selection process (Lancer et al. 2019). Figure 8 shows the equivalent BSTMD2 raster plots (n = 6). BSTMD2 responses to the TO case generally lasted for the entire stimulus duration, indicative of its large receptive field across both eyes (Figure 8A). In the background alone case, BSTMD2 exhibited strongly directional responses (Figure 8B) with intermittent excitation to leftward motion (BSL) and more consistent strong inhibition to rightward motion (BSR). In the background-with-target trials (BSLT, BSRT) target responses are reduced, especially in the rightward (BSRT) case (Figure 8C). Here we see further evidence of selection-like processes, with initial strong inhibition (to background features) frequently replaced by strong excitation (to the target). These inhibition to excitation transitions do not always occur at the same time point indicating that this switch is not simply due to the target reaching a specific receptive field location. This is consistent with previous descriptions of selective attention, where switches can occur at varying times (Wiederman & O’Carroll 2013). In the BSLT trials we also see reduced responses. This is despite leftward moving backgrounds still increasing excitation when presented alone. While this could potentially be due to longer range spatial inhibition between the target and nearby background features, the inconsistent time course of breakthroughs to higher response levels are more indicative of prior attention to low-salience background features subsequently out-competing the higher salience target.

**Figure 8:**
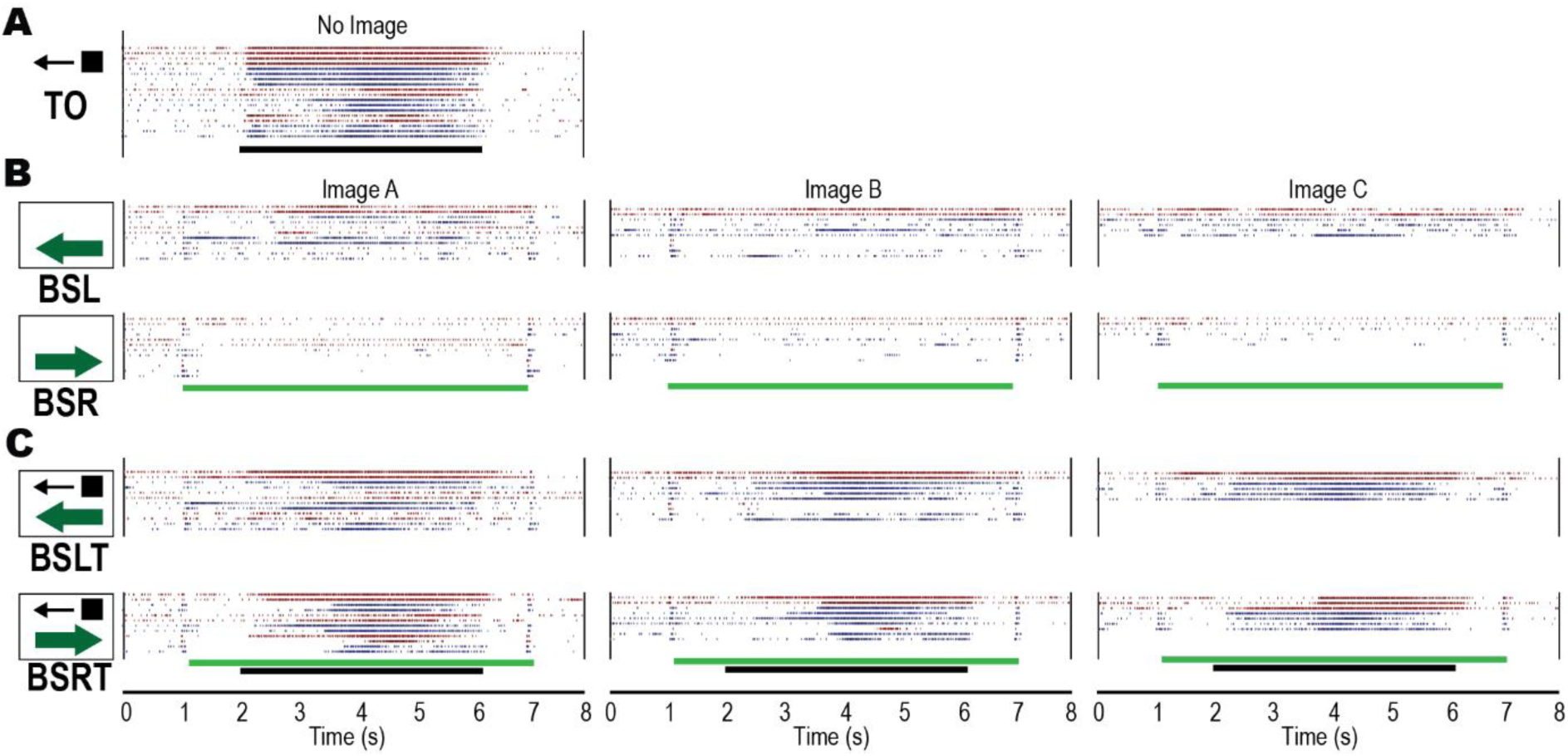
As observed in CSTMD1, BSTMD2 raster plots reveal that moving backgrounds cause target responses to become less sustained and consistent. **A,** Target only responses are robust in the preferred direction. **B,** Backgrounds moving in the preferred direction produce many more ‘false-positive’ responses than CSTMD1. Background features moved in the anti-preferred direction induce inhibition. **C,** Background and target moved in the preferred direction result in spike activity from different parts of the receptive field. Background and target moved in opposite directions result in distinct target or background responses.

Previous experiments and modelling suggest that most natural images contain features that can elicit responses in STMD neurons, although not as robustly as embedded targets (Wiederman & O’Carroll 2011). Here very strong responses to background features were rare, however our patterns were moved at a relatively slow velocity (15°/s) compared with the known preference of the neurons for targets moving around 5-6 times faster than this (Geurten et al. 2008, Wiederman). Nevertheless, we found several examples where background features robustly stimulated the neuron, allowing us to examine the fine structure of the response time course in more detail. Figure 9 (A-J) shows traces for individual examples of such responses either for the background alone (BO, green lines), target alone (red lines) or B+T (black lines). In all these examples the only difference is the presence or absence of the target or background. All other parameters (such as image and image phase) are the same. Throughout these examples, an asterisk (*) marks periods when the B+T response is similar to responses to background-only trials, suggesting that at these time points, the neuron is selecting the background feature, rather than the target,

**Figure 9:**
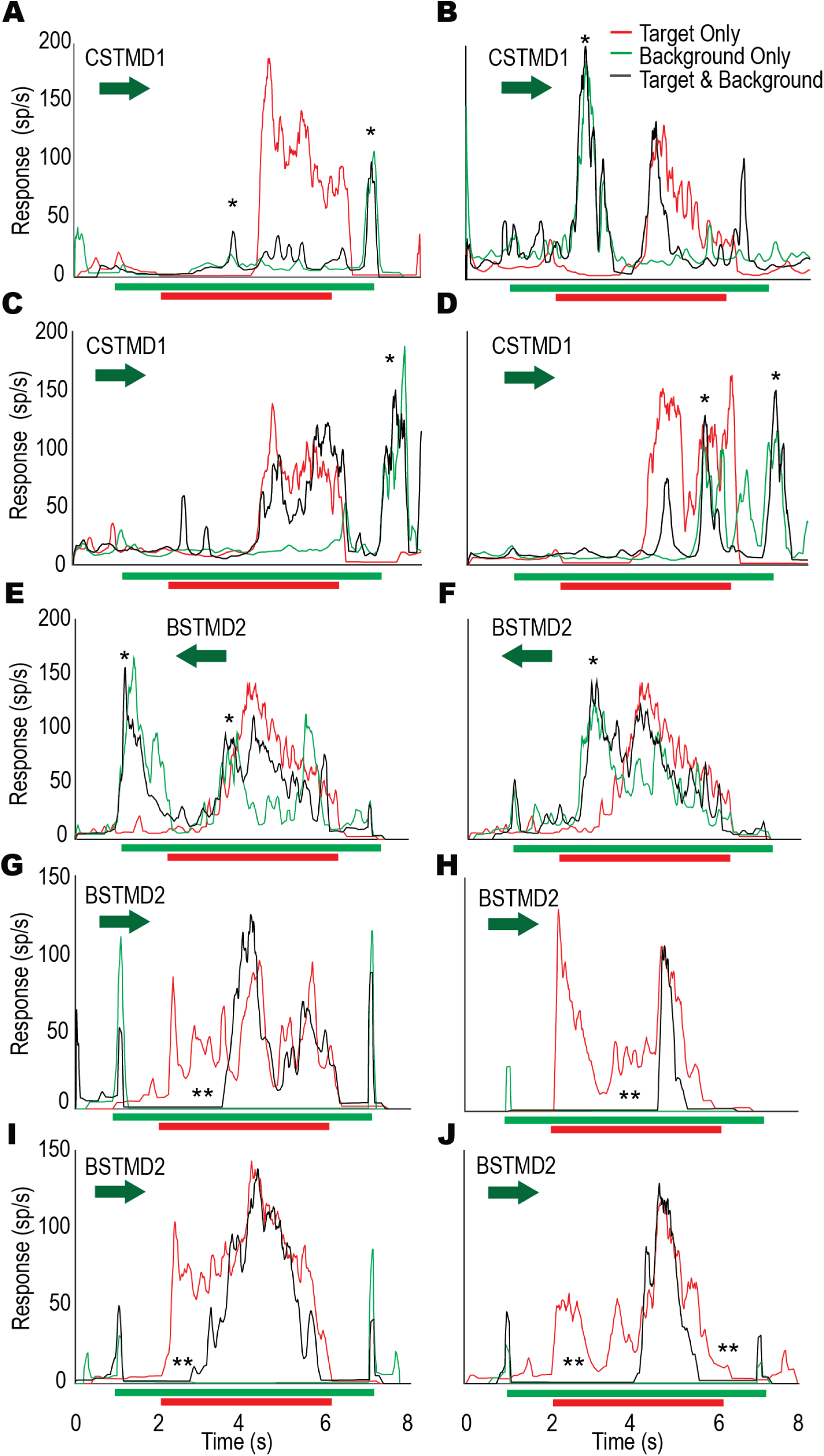
Individual examples comparing target-only (red), background-only (green) and target with background trials (black). Stimulus timings marked below each figure. **A)** An example in CSTMD1 where when the background and target are presented, the responses match the background-alone trial which reveals that sometimes a background feature can win the selection process. **B-D)** Examples from three neurons where the background generates strong responses and causes some suppression of the target response. **E-F)** Examples from BSTMD2 where the neuron switches between background and target features. **G-J)** BSTMD2 trials exhibit numerous examples of the neuron being inhibited by background motion despite target features, including switches between background and foreground features.

Figure 9A shows an example where introduction of the background completely suppresses CSTMD1’s response to the target (compare the black and red lines) while maintaining a characteristic response to the background features, both prior to and subsequent to the target’s appearance. Figures 9B and 9C show examples where both the target and a background feature generate periods with strong excitatory responses resembling those for either stimulus alone, indicating that the system can switch from one to the other (though only for a brief response to the target in 9B). Figure 9D shows an example where the response to both background and target (B+T, black) remains better matched to the background only (green) than target only (red), even when the target is within the most sensitive part of the receptive field. This example illustrates how the intermittent responses seen in the raster plots (Figure 7 and Figure 8) can arise. While the first half of the trial (2-4s) exhibits very weak responses, the second half (4s onwards) produces intermittent responses which are well matched to background features, including at ∼7s where the target is no longer present.

Figure 9E and Figure 9F show two example responses from a single BSTMD2 neuron, with the only difference being the starting position (phase) of the background image. In both trials, responses transition from the background features to the target mid-trial. Interestingly, in both trials the B+T responses (black line) is slightly weaker than in the target alone case (red line). This is unlike our previous descriptions of selective attention in CSTMD1, where once selected the response continues as if the distractor does not exist. Therefore, in BSTMD2 there may be an additional inhibitory effect of background features, perhaps related to its broader size selectivity (Figure 2E).

Figures 9G-J show several examples from BSTMD2 where the background is moving in a direction opposed to the target. In these trials the background-alone response is always inhibitory (green line) but this inhibition also ‘wins’ over the excitatory target response (red line) for varying periods when the background is presented with a target (black line). Double asterisks (**) mark times when an inhibitory background feature (green line) is selected over an excitatory target (red line).

### Selection for Faster Velocities

We have so far shown that even features within slow-moving backgrounds can be the subject of selection. How might a faster-moving background, as might be experienced during quick turns towards targets during pursuit flights, affect neuronal responses to a fast or a slow-moving target? To investigate this, we chose a single, cluttered background image (Image A, Figure 5A) and recorded responses in CSTMD1 to targets moving at one of three velocities (15, 35 and 90°/s) whilst the background moved in one of three different velocities (15, 35 and 90°/s) in the neuron’s preferred direction (rightwards). We measured spiking activity in an analysis window corresponding to the moving target. As faster target velocities evoke responses over a shorter duration of time, we changed our analysis window (Figure 10A) to correspond to a period when the target moved over a constant region of space (50°) within the receptive field. We used these same analysis windows even when the target was not present.

**Figure 10:**
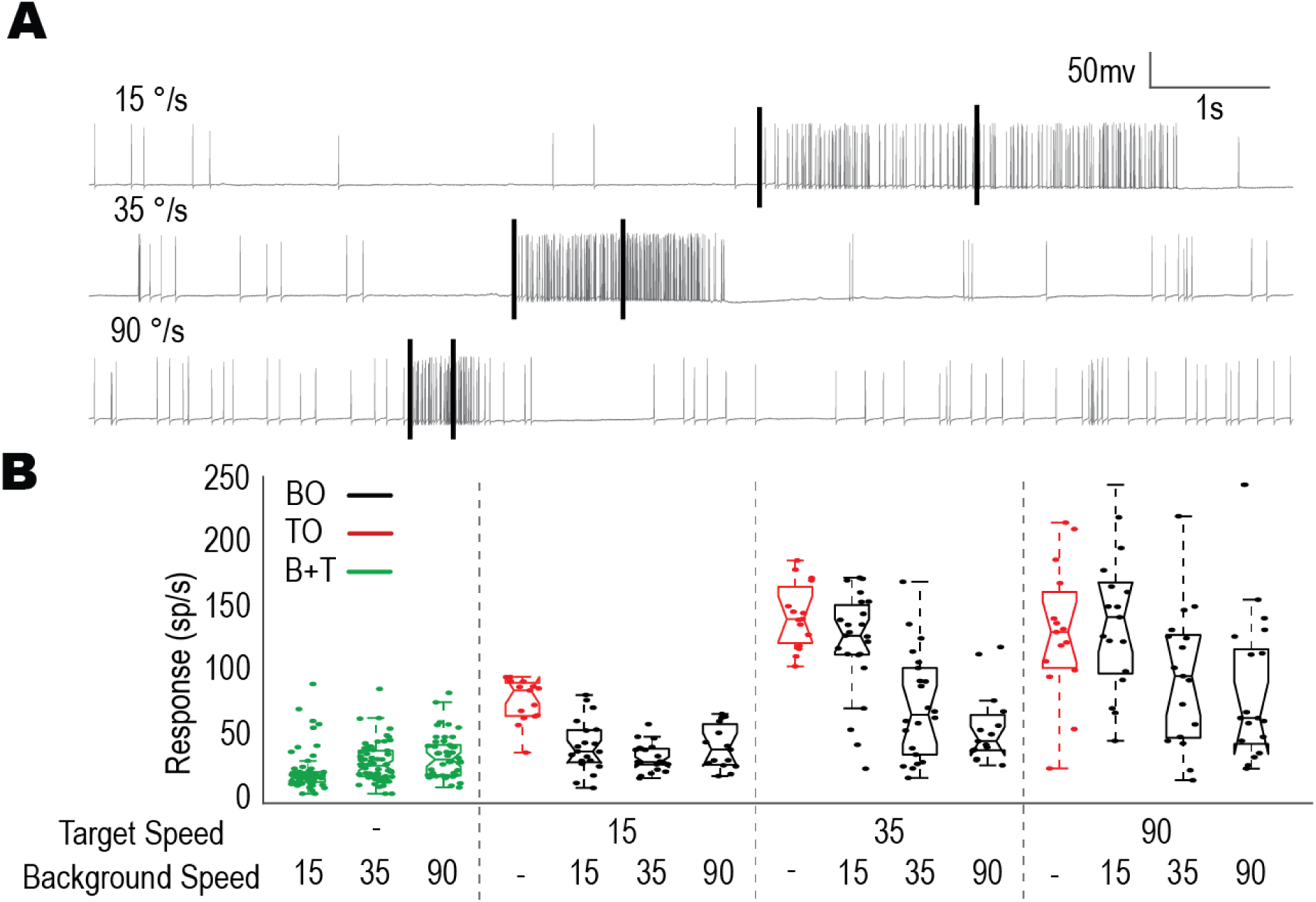
Velocity influences the selection of moving targets. **A,** Three raw traces taken from targets of varying velocities showing the difference in response and time-course. Black bars indicate an analysis window of equal size (50°). **B,** Box plot showing CSTMD1 responses over the analysis window, for the image conditions (target only -red, background only -green, target and background -black) across combinations of target and background velocities. This includes three target-only velocities, three background only velocities and nine target speed and background speed combinations (all in the preferred direction). Target response is more robust when the target moves as fast, or faster than the background.

Figure 10B shows the spike-rate from these windows for each of fifteen conditions: 3 target-only controls for the 3 velocities (red); 3 corresponding background-only controls (green); and 9 background/target speed combinations (black). The target-only trials exhibited the expected velocity tuning as previously described (Dunbier et al., 2012) with stronger responses for the two higher velocities. Although higher speeds did also produce slightly stronger responses in background-only trials, these were still very weak at all speeds compared with even the slowest target-only condition.

B+T trials showed two important interactions: Firstly, responses continued to increase at the target speed increased, consistent with the inherent velocity tuning. However neuronal responses for each target speed then decreased as the background speed increased, most likely reflecting for frequent switches of attention to the (now more salient) background features. Both these factors had a statistically significant effect on neuronal response (two-way ANOVA, n=11; background speed, p < 0.0001; target speed, p < 0.0001). To de-confound the effects of velocity and feature selection on spike-rate, we normalized the target-with-background trial responses by dividing these responses by the mean response of the corresponding target-only trials (for example all background-with-target trials with the target moving 90°/s had their responses divided by the mean of the 90°/s target-only trials). With this normalization, the two relationships were maintained (two-way ANOVA, n=11; background speed, p < 0.0001; target speed, p = 0.0004). This analysis reveals that higher velocity targets not only generate stronger responses, but also are more likely to be selected.

How does this interaction occur? If selective attention underlies the interactions between background and target, the data should show that faster target speeds and slower background speeds increase the likelihood of target selection. In all of the B+T conditions, there are responses that match the corresponding target-only distribution and others that match the background-only distribution, with relatively few points half-way in between. This ‘selection’ is particularly visible in the B+T trials with a background speed of 90°/s (3^rd^ box-plot of each set, Figure 10B). All three show a marked division between responses around 50 spikes/s and response greater than 100 spikes/s. In the slow-target case (15°/s) all responses are clustered around 50 spikes/s. In the intermediate speed case (35°/s) most trials match the slow target, but with two outliers greater than 100 spikes/s. In the fast-target case (90°/s) there is still a cluster of responses centred around 50 spikes/s but now there are many more (7 total) outliers above 100 spikes/s. Not all conditions exhibited as strongly a bimodal distribution as the 90°/s-background cases, however switching may explain these intermediate cases.

The B+T trials demonstrate that both target and background speed have an important effect on the selection process of CSTMD1. Both relationships indicate that velocity is a key determinant in selection. Features better matched to CSTMD1’s preferred velocities (i.e. either 90°/s targets or background features) are preferentially selected over slower features, even when other aspects of those features (contrast, size etc) are lacking.

### Modelling Selective Attention

In an effort to capture the selection between targets and background features observed in the individual examples in CSTMD1 and BSTMD2 (Figure 9), we developed five models of neuronal interactions to compare with the physiological data, a similar approach to our previous investigation of selective attention within the excitatory receptive field (Wiederman & O’Carroll 2013). The models were designed to predict the background with target responses (BSLT, BSRT) from different combinations of the target-alone (TO) and background-alone (BSL, BSR) physiological data (Figure 11A). For each animal (CSTMD1, n=12; BSTMD2, n = 6), we first found the mean response of the TO trials to represent the Target-Only input. For the Background-Only input, we separated trials based on the image used, the background direction and the background starting location (phase) and then found the corresponding mean. This resulted in a total of twelve different Background-Only inputs (three images, two phases, two directions). Both the mean TO and set of mean BSL/BSR responses were then segmented into 5 ms bins and combined according to the rules of each model variant (5 variants, details below). We then calculated an error between model output and the physiological B+T data (matched to the corresponding image, phase and direction).

**Figure 11:**
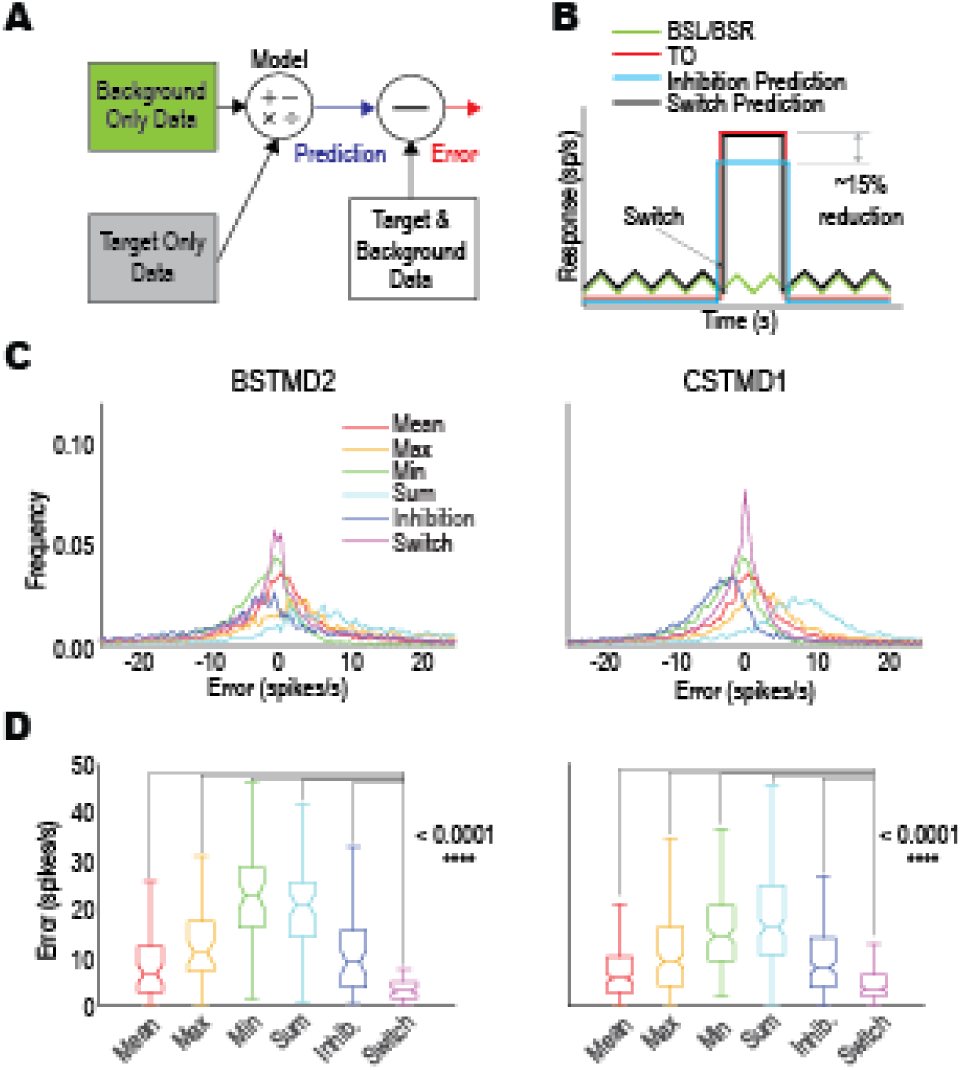
A selective attention model that includes the ability to switch, outperforms other model variants. **A,** A schematic of the modelling approach. The target-only (TO) and background-only (BSL, BSR) data are used to predict a background with target response and this is then compared to the background with target data (mean error). **B,** Illustration of a model variant; Switching (black line) generated from the corresponding TO (red line) and background-alone (green line) responses. The error is the difference between the model output and the physiological background and target responses (not shown). **C,** Histogram of errors for the model variants Models include Mean (average of background-only and target-only responses), Max (maximum of background-only and target-only responses), Min (minimum of background-only and target-only responses), Sum (combination of background-only and target-only response and Switch (smaller error between target-only and background-only responses). **D,** Boxplots showing mean error across each trial for each model variant. A smaller value represents a better match between model and data. The Switch model is the best match for both CSTMD1 and BSTMD2 (p < 0.0001, t-Test, Bonferroni correction).

The six variant models tested are as follows: 1) ***Mean***: the average of the Target-Only or Background-Only mean responses (each in 5 ms bins), 2) ***Max***: the maximum of the Target-Only or Background-Only mean responses, 3) ***Min***: the minimum of the Target-Only or Background-Only mean responses, 4) ***Sum***: the addition of the minimum of the Target-Only or Background-Only mean responses, 5) ***Switch***: the smaller of the errors between either the Target-Only or Background-Only mean responses (Figure 11B).

A distribution for the errors across all model variants is shown in Figure 11C. The *Switch* model resulted in the smallest errors in both CSTMD1 and BSTMD2 indicated by the high peaks at zero and narrow flanks. The *Sum* and *Max* models generated large over-estimates of the response while the *Min* model produced underestimates. We further calculated the mean absolute error for each trial (averaged across 5 ms bins) and treated each trial as an independent error (Figure 11D). The *Switch* model produced the smallest mean error of the six variants tested. The Switch model was significantly better than all other variants (p < 0.0001 T-Test with Boneferroni correction, in all five comparisons). This further supports that background interactions with these two large-field STMD neurons functions through a competitive attention mechanism that switches between background features and the target.

### Detection Error Trade-Off Performance

In behavioural contexts, eventually all detection tasks result in a simple binary decision: in this case that would be to initiate pursuit, or not? How (and whether) STMDs are involved in this behaviour remains unknown, but assuming they play a role, how might the responses of STMDs in visual clutter affect this kind of decision making? While relating the spike-rate of these neurons directly to these behaviours remains infeasible, it is still possible to examine how clutter might affect the detection capability of STMDs towards these ends. At what point is a known salient target indistinguishable from a moving background feature?

To examine this question, we applied a “detection error tradeoff” (DET) analysis applied to an extended set of our moving background experiment (as in Figure 6) in which the target contrast was also varied. If we assume that the system is designed to detect small targets (such as the one we present) while ignoring background features, there are several logical inferences which can be made from these trials. For example, in background-only trials, there is no ‘true’ target and thus any strong response can be considered to be a false-positive (FP). Likewise, when a salient target is presented alone with no neuronal response, this represents a false negative (FN). When both target and background are presented it is impossible to categorically distinguish between target and background responses and thus positive responses remain ambiguous, however the lack of a response when a target is present is also a FN. A summary of this logic is shown in Figure 12A. A useful metric that can be derived from this data is the Detection Error Trade-Off (DET), which compares FP and FN events and is a measure of tracking performance (Martin et al. 1997). The advantage of the DET, is that it does not assume a single fixed threshold for detection but instead shows how changes in assumed threshold affect the prevalence of FP and FN events.

**Figure 12:**
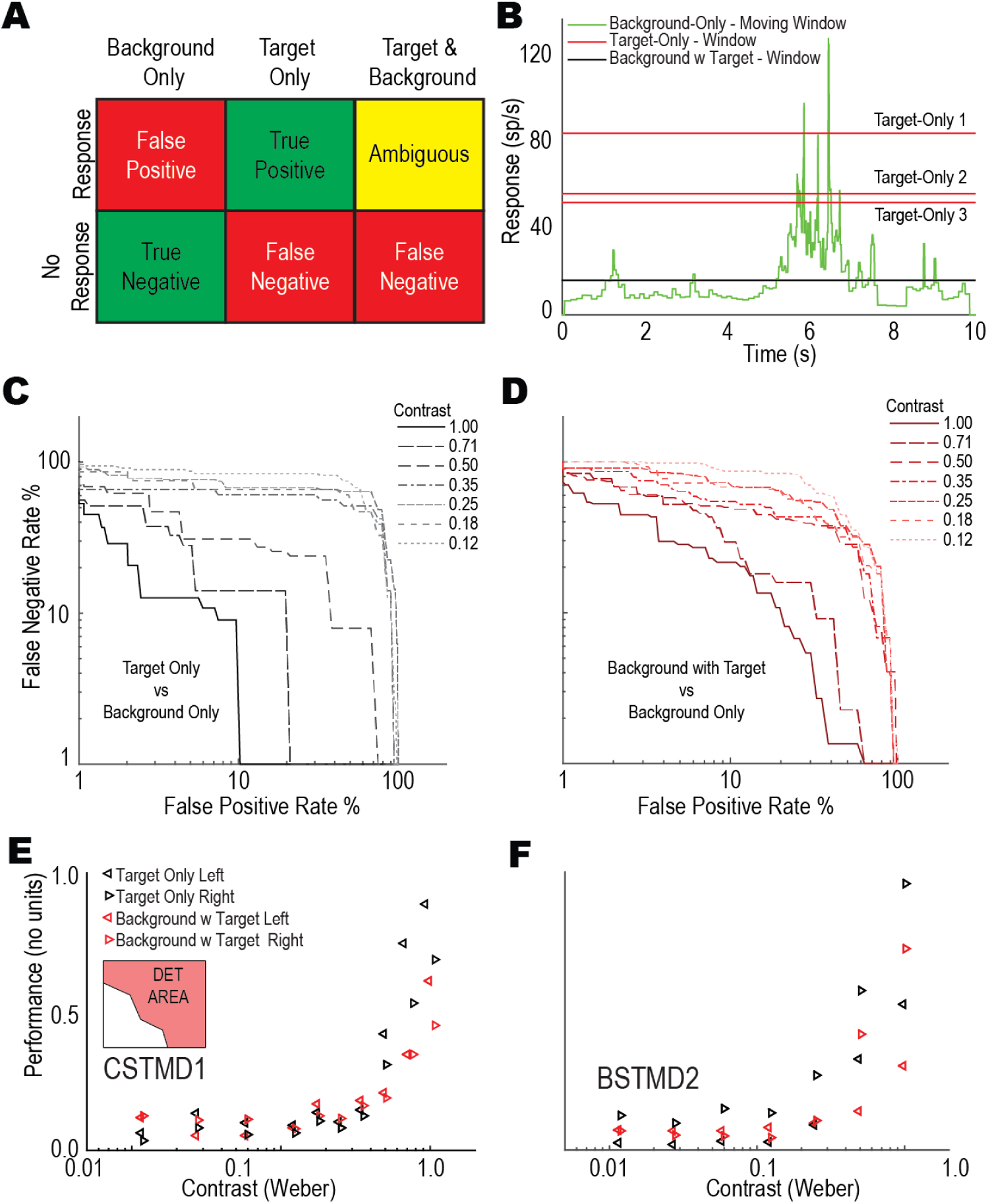
Performance of STMD neurons reduced in clutter regardless of detection threshold selection. **A,** Table showing deductions that can be made based on a threshold-based response mechanism. Strong responses in target with background trials have an ambiguous interpretation. **B,** Example of a rolling-window measuring background response over the course of an entire background-only trial (green line) with the windowed responses of target-only controls and background with target responses. Regardless of chosen response threshold, it is impossible to achieve zero False Positives and zero False Negatives simultaneously. **C,** Detection Error Tradeoff curves for CSTMD1 for all measured contrasts to target-only trials. **D,** Detection Error Tradeoff curves for CSTMD1 for all measured contrasts to background with target trials. Background with target trials show worse performance than target-only trials regardless of threshold selection. **E, F,** Area above logarithmic Detection Error Tradeoff curve for all contrasts separating left and rightwards moving backgrounds for CSTMD1 (E) and BSTMD2 (F). Both neurons show more reliable performance when the background moves against the preferred direction of the STMD.

To measure the DET rate, we established the definitions of FP and FN events. For this analysis we separated target-only trials and background-with-target trials into two separate categories and performed the following analysis on both datasets. To calculate the FN rate, we used a 500 ms window for each trial taken from the most sensitive region of the receptive field (adjacent to the midline). Trials that failed to meet a (variable) threshold were classed as FN events. FP rate was determined by aggregating the background-only trials. As these trials contained no target, we calculated windows along the length of the entire trial. Using a sliding window method (500 ms) we generated an instantaneous snapshot of the background trials at each time point (Figure 12B – green line). Using this aggregated data, we then determined a FP-rate for any given threshold by calculating what percentage of each trial was above threshold. To calculate the DET curve for target-only trials, we used a variable threshold simultaneously calculating the FN-rate from the target-only trials and the FP-rate from the background-only trials. An example of how this process works is shown in Figure 12B. Here there were three target-only controls and a single B+T example. If one changes the theoretical threshold from zero spikes/s up through 120 spikes/s, the total FP rate (the proportion of the green line exceeding the threshold) declines, while the FN rate would increase as the threshold exceeds the target-only controls. Target-only trials are shown in Figure 12C and B+T responses in Figure 12D (CSTMD1 responses across all images and background directions). We observe a performance reduction as the contrast of the target is reduced (the upper right of this plot represents larger probabilities of FP and FN events). Even at the highest contrast, it is impossible to achieve both low FP and FN rates, indicative of the background images occasionally producing strong target-like responses when presented alone. This also demonstrates that the introduction of clutter increases the rate of FP and FN events (compare Figure 12C and Figure 12D).

We calculated the area above the curve (DET area) and normalized it such that a value of 1 represented 0% FP and 0% FN, while 0 represented 100% FP and 100% FN. The resulting DET Areas (separated by background direction) are plotted in Figure 12E (CSTMD1) and Figure 12F (BSTMD2). The DET Area shows that lower target contrast reduces the DET performance and that the presence of clutter also always reduces the DET performance (red is uniformly lower than black at high contrast). This difference due to clutter is further evidence for a target-clutter interaction. In the target-only/background-only comparison, the differences seen are due to the relative activity of the two conditions (i.e. low contrast targets are more likely to respond less strongly than intermittent clutter features). However, in the clutter-with-target/background-only comparison, the weaker overall response is due to the direct inhibitory effects caused by the background on the target responses (i.e. the target/background interaction). Finally, there is a directional difference in both cell types: both show poorer performance when the background moves in the preferred direction. This is best explained by a much higher false-positive rate introduced by preferred-motion features in the background. This indicates that when in the presence of moving clutter, both CSTMD1 and BSTMD2 are more reliable target detectors when there is opposing motion (rather than parallel motion) within a scene.

## Discussion

The presence of background clutter in a scene confounds target detection by changing local contrast as well as by presenting distracting features. We utilised the grey strip paradigm to remove changes in target contrast, thus isolating long-range effects to those due to optic-flow and competition with other background features. In other insects, background motion may completely suppress neuronal responses, as observed in descending neurons (Nicholas et al. 2018) and some lobula motion detectors (Keles et al. 2020, Stadele et al. 2020). Here we show that dragonfly STMD responses exhibit trial-by-trial variability, which our modelling suggests is due to selective attention rather than surround suppression. In any individual trial, STMD responses can be either unaffected by the presence of clutter or selectively respond to background features (excitatory or even inhibitory) rather than the high salience target.

Single targets presented simultaneously in both visual hemispheres provide evidence that selective attention operates across the brain. This selection can represent either excitatory or inhibitory stimuli and is clearly biased by a preceding ‘primer’. As both CSTMD1 and BSTMD2 can select inhibitory stimuli, the underlying selective processing is likely to occur outside of either neuron, suggesting the involvement of a network of higher-order neurons. Each hemisphere may possess an independent bottom-up selection network, with the final determination between hemispheres occurring at a higher level and perhaps involving neurons like CSTMD1, which have large diameter axons that cross between the left and right optic lobes. Whether two neurons with overlapping receptive fields are selecting the same target simultaneously (i.e. centralised selection) remains to be determined, although our observation that attentional switches between foreground and background features are more common in BSTMD2 than in CSTMD1 suggests that there may be more than one attentional network operating in parallel.

Our previous research described that robust responses to background features were rare (Wiederman & O’Carroll 2011), as few background features strongly matched the finely tuned selectivity of STMD neurons. Although this result was replicated in the present study, we did find numerous individual examples of both CSTMD1 and BSTMD2 responding to background features even when a high-salience target was present. This echoes recent findings where low-contrast targets can maintain selection (‘lock-on’) despite the introduction of high-contrast distracters (Lancer et al. 2019). Our backgrounds were presented prior to target appearance, a priming that may have biased the selection of background features. This scenario is more representative of the real-world ecology of hawking dragonflies, where a moving background is often present. The ability of target-detection pathways to select weak or inhibitory background features within moving backgrounds raises new questions about how target pursuit behaviours emerge from these underlying neuronal networks.

In the selective attention process, how is the winner determined? Our clutter experiments reveal that overall spike activity is not the determining factor. If it were, we might expect robust responses to clutter features, rivalling those of the highly salient target. Such an effect is observed in primate cortical cells where selection can be biased by ‘enhancing’ the weak stimulus until it is the winner (Martinez-Trujillo and Treue, 2002; Reynolds and Desimone, 2003). Instead we sometimes observe weak responses well matched to background features despite the presence of a target which when presented alone produces a strikingly strong response.

It is likely that the biasing of subsequent selection by primers that we observe is mediated by a form of ‘predictive gain modulation’ where individual target responses are increased when on predictable forward trajectories, whilst other parts of the receptive field are suppressed (Wiederman & Fabian et al. 2017, Fabian et al. 2019). Similar temporal cuing behaviour is also observed in flies where attention can be biased by preceding cues (Sareen et al. 2011, Koenig et al. 2016). Although primed and selected background features may not exhibit strong STMD responses, the concomitant surround suppression may be sufficient to suppress distracters (enabling the ‘lock-on’ effect described by Lancer et al. 2019). Such processing would effectively inhibit ‘switches’ between targets, which are more rarely observed in physiological recordings. However, this does not explain individual trials where the target response is selected, but only transiently. In these examples, muted target response sometimes lasted only briefly. These switches are away from the highly salient target even though it has already been within the visual field for seconds and could potentially have already facilitated responses of local neurons in lower-order networks. Further investigation is required to understand these responses and how they may relate to regulation by the overall neuronal network.

We have shown that velocity is an important criterion for the selection of features, with those matched to optimal STMD velocity tuning being selected preferentially to slower moving features. This result agrees with experiments examining predictive gain modulation where velocity is a key determinant of facilitation strength (Fabian et al. 2019). Features moving at optimal velocities (∼90 deg/s) may facilitate their salience, thus improving their chance for selection. However, it is important to note, we sometimes observed switches to clutter features despite the more slowly moving background.

While we previously showed that relative motion is not required for target detection (Wiederman & O’Carroll 2011), here we described that relative motion reduces the likelihood of robust detection, even if the background motion is opposite to that of the target. Additionally, relative motion did not increase neuronal responses, even when the target was detected. This is unlike in humans where relative motion is an important cue for detection (Smeets & Brenner 1994). In BSTMD2 trials, although relative motion did not increase spiking activity, the overall difference is increased in trials of opposed background motion. In these examples, background motion in the anti-preferred direction reduces spike-rate below spontaneous levels, so when a target appears it can elicit a larger response change (i.e. switching from inhibition to excitation). Our results reveal that the most robust target detection occurs when background motion is minimized, which may match dragonfly hunting behaviour (Bomphrey et al. 2016) and the saccadic flight dynamics of flies (Tammero & Dickinson 2002). Our experiments were conducted in open-loop with a restrained animal. In contrast, closed-loop experiments in flies reveals efference copies from motor commands (Kim et al. 2015, Kim et al. 2017) subtract the fly’s intended movement from its widefield motion detection pathway via modulating feedback. We cannot rule out a similar effect existing in the dragonfly target detection pathway, potentially enhancing responses to targets moving against a moving background via relative motion cues.

If facilitation and a near optimal velocity is not always sufficient to win the competitive selection and there are no relative motion cues from widefield motion (Evans et al. 2019), then what drives the selection process? One explanation is that the mechanisms determining selection are not only based on the individual STMD’s tuning for that property (e.g. size, velocity). It is possible that neurons with different tuning to larger or slower features feed into a bottom-up selective attention process, permitting switches to these less optimal stimuli. Dragonflies have been shown to minimize the relative ‘slip’ of a target, aiming for no relative target velocity across the eye (Mischiati & Lin 2014). In these scenarios, STMDs should display minimal activity, except to the background features induced by ego-motion. How might this apparent contradiction be reconciled? In close-loop pursuit, STMDs would respond robustly to target ‘slippage’ from the midline, effectively encoding the error signal in pursuit. This matches CSTMD1’s preference for targets moving away from the midline. If STMDs are used in this capacity, the ability of the selective attention system to prioritize low-salience targets would be necessary to maintain tracking.

Alternatively, the role for these STMDs may be limited to the detection phase whilst hawking, with only a minor contribution in the subsequent pursuit phase of any engagement. Our findings match the hawking behaviours of dragonflies like *Hemicordulia*, which often attempt to hover in regions where the background is clear sky. These situations maximize the contrast of potential prey and minimize the interference of motion from background distracters.

Despite these limitations, STMDs still respond robustly to targets in challenging dynamic, cluttered scenarios. Dragonflies do not always choose their engagements, with territorial or sexual conspecific encounters often commencing with the target hidden in front of cluttered background. In these circumstances, and the complex pursuit flights that follow, STMDs are still capable of generating robust responses to moving targets whilst suppressing the responses of distracting clutter features. This is at odds with findings from flies, which appear to suppress target detection in the presence of moving clutter (Keles et al. 2020), perhaps making dragonflies better suited to dynamic and complex predatory pursuits.

## Acknowledgements

This research was supported by the Australian Research Council’s Future Fellowship scheme (FF180100466) and the Swedish Research Council (VR 2018-03452). We thank the manager of the Adelaide Botanic Gardens for allowing insect collection and behavioural recordings.

